# Delineating the molecular and phenotypic spectrum of the *SETD1B*-related syndrome

**DOI:** 10.1101/2021.02.11.430742

**Authors:** Marjolein J.A. Weerts, Kristina Lanko, Francisco J. Guzmán-Vega, Adam Jackson, Reshmi Ramakrishnan, Kelly J. Cardona-Londoño, Karla A. Peña-Guerra, Yolande van Bever, Barbara W. van Paassen, Anneke Kievit, Marjon van Slegtenhorst, Nicholas M. Allen, Caroline M. Kehoe, Hannah K. Robinson, Lewis Pang, Selina H. Banu, Mashaya Zaman, Stephanie Efthymiou, Henry Houlden, Irma Järvelä, Leena Lauronen, Tuomo Määttä, Isabelle Schrauwen, Suzanne M Leal, Claudia A.L Ruivenkamp, Daniela Q.C.M. Barge-Schaapveld, Cacha M.P.C.D. Peeters-Scholte, Hamid Galehdari, Neda Mazaheri, Sanjay M Sisodiya, Victoria Harrison, Angela Sun, Jenny Thies, Luis Alberto Pedroza, Yana Lara-Taranchenko, Ivan K. Chinn, James R. Lupski, Alexandra Garza-Flores, Jefferey McGlothlin, Lin Yang, Shaoping Huang, Xiaodong Wang, Tamison Jewett, Gretchen Rosso, Xi Lin, Shehla Mohammed, J. Lawrence Merritt, Ghayda M. Mirzaa, Andrew E. Timms, Joshua Scheck, Mariet Elting, Abeltje M. Polstra, Lauren Schenck, Maura R.Z. Ruzhnikov, Annalisa Vetro, Martino Montomoli, Renzo Guerrini, Daniel C. Koboldt, Theresa Mihalic Mosher, Matthew T. Pastore, Kim L. McBride, Jing Peng, Zou Pan, Marjolein Willemsen, Susanne Koning, Peter D. Turnpenny, Bert B.A. de Vries, Christian Gilissen, Rolph Pfundt, Melissa Lees, Stephen R. Braddock, Kara C. Klemp, Fleur Vansenne, Marielle van Gijn, Catherine Quindipan, Matthew A. Deardorff, J. Austin Hamm, Abbey M. Putnam, Rebecca Baud, Laurence Walsh, Sally A. Lynch, Julia Baptista, Richard E. Person, Kristin G. Monaghan, Amy Crunk, Jennifer Keller-Ramey, Adi Reich, Houda Zghal Elloumi, Marielle Alders, Jennifer Kerkhof, Haley McConkey, Sadegheh Haghshenas, Genomics England Research Consortium, Reza Maroofian, Bekim Sadikovic, Siddharth Banka, Stefan T. Arold, Tahsin Stefan Barakat

## Abstract

Pathogenic variants in *SETD1B* have been associated with a syndromic neurodevelopmental disorder including intellectual disability, language delay and seizures. To date, clinical features have been described for eleven patients with (likely) pathogenic *SETD1B* sequence variants. We perform an in-depth clinical characterization of a cohort of 36 unpublished individuals with *SETD1B* sequence variants, describing their molecular and phenotypic spectrum. Selected variants were functionally tested using *in vitro* and genome-wide methylation assays. Our data present evidence for a loss-of-function mechanism of *SETD1B* variants, resulting in a core clinical phenotype of global developmental delay, language delay including regression, intellectual disability, autism and other behavioral issues, and variable epilepsy phenotypes. Developmental delay appeared to precede seizure onset, suggesting *SETD1B* dysfunction impacts physiological neurodevelopment even in the absence of epileptic activity. Interestingly, males are significantly overrepresented and more severely affected, and we speculate that sex-linked traits could affect susceptibility to penetrance and the clinical spectrum of *SETD1B* variants. Finally, despite the possibility of non-redundant contributions of *SETD1B* and its paralogue SETD1A to epigenetic control, the clinical phenotypes of the related disorders share many similarities, indicating that elucidating shared and divergent downstream targets of both genes will help to understand the mechanism leading to the neurobehavioral phenotypes. Insights from this extensive cohort will facilitate the counseling regarding the molecular and phenotypic landscape of newly diagnosed patients with the *SETD1B*-related syndrome.

## INTRODUCTION

*SETD1B* encodes a lysine-specific histone methyltransferase that methylates histone H3 at position lysine-4 (H3K4me1, H3K4me2, H3K4me3) as part of a multi-subunit complex known as COMPASS^1,2^. The *SETD1B* protein consists of 1966 amino acids and has several (presumed) functional domains (Figure 1). The N-terminus contains an RNA recognition motif (RRM), whereas the middle region is characterized by two long disordered regions that differ from other homologs^3,4^, a conserved lysine-serine-aspartic acid^5^ (LSD) motif and a coiled coil structure. At the C-terminus, *SETD1B* harbors a catalytic SET domain crucial for histone methyltransferase activity, bordered proximally by the N-SET domain including a conserved^6^ WDR5-interacting (WIN) motif, and distally by the post-SET domain. H3K4me3 is enriched at promotor and transcription start sites whereas H3K4me1 and H3K4me2 are enriched at enhancer sites, therefore^7^ being associated with active gene transcription and euchromatin. Indeed epigenetic changes have been observed in both animal models and patient material^8–10^ at promoters and intergenic regions, confirming that *SETD1B* epigenetically controls gene expression and chromatin state. In addition, *SETD1B* is^11^ constrained for both missense and loss of function variants.

**Figure 1:**
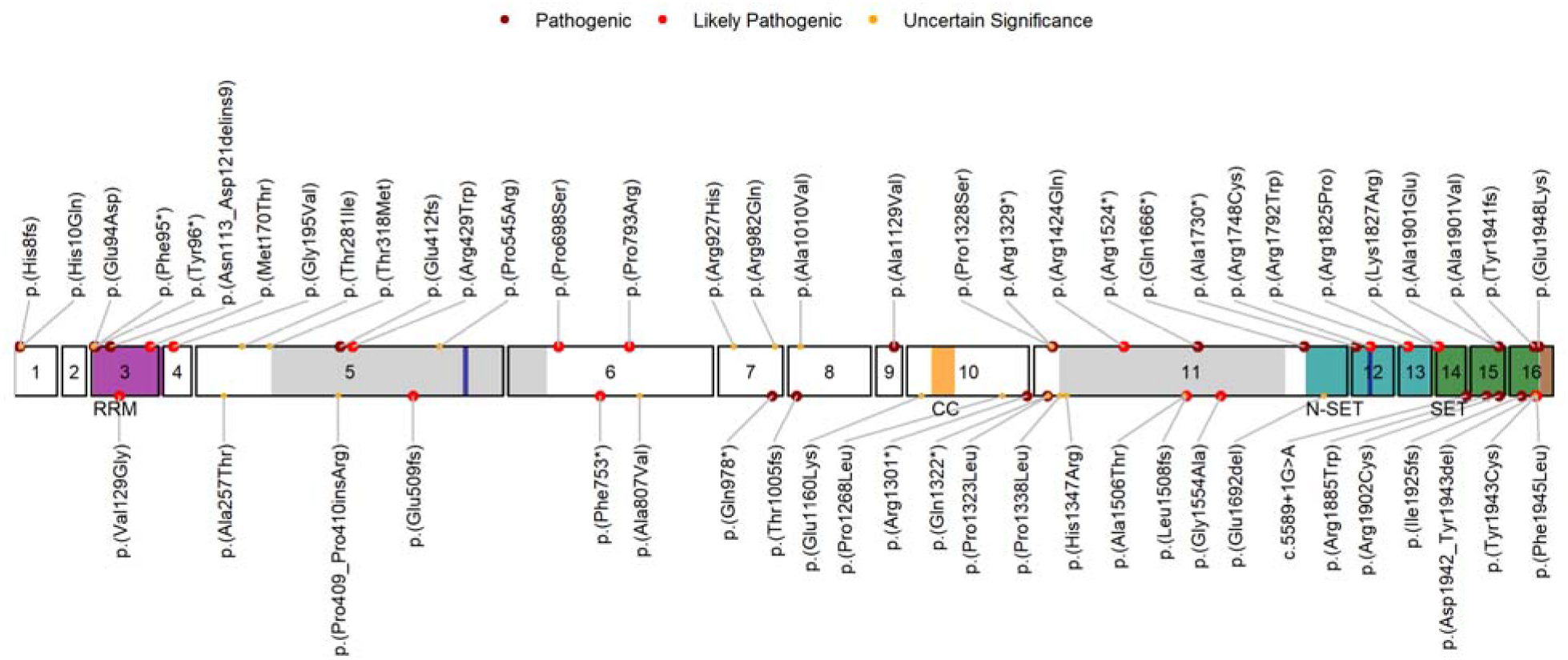
Schematic representation of *SETD1B* variants in this study cohort (major circles, top labels) and in literature (minor circles, bottom labels). The RRM (residues 94-182), coiled-coil (CC) (residues 1173-1204), N-SET (residues 1668-1821), SET (residues 1822-1948) and post-SET (residues 1949-1966) domains in respectively magenta, orange, cyan, green and brown, the largely disordered regions (residues 320-682 and residues 1338-1640) in light grey, and the LSD (exon 5, residues 577-583) and WIN (exon 12, residues 1745-1750, within N-SET) motifs both in blue. Will be available upon peer review

Consistent with this, pathogenic variants in *SETD1B* have been associated with a syndromic intellectual developmental disorder including seizures and language delay (IDDSELD, OMIM #619000). To date, clinical features have been described for eleven affected individuals with (likely) pathogenic *SETD1B* sequence variants^8,12–15^. Individuals with microdeletions encompassing *SETD1B* have also been described^8,16–19^, however most of these deletions encompass additional genes making phenotypic comparisons challenging. In this study, we further delineate the clinical phenotype associated with *SETD1B* sequence variants, by describing 36 additional individuals. Comparing these new cases to the published ones provides a comprehensive molecular and clinical characterization of the *SETD1B*-related syndrome. In addition, using protein modelling, *in vitro* assays, and genome-wide methylation signatures we investigate the effects of selected variants, providing further evidence for their pathogenicity. Together, this work expands the molecular and phenotypic landscape associated with *SETD1B* variants.

## METHODS

### Ethics statement and cohort inclusion

After identification of three individuals with *SETD1B* variants at Clinical Genetics of Erasmus Medical Center, additional cases were identified using GeneMatcher^20^, the Dutch Datasharing Initiative^21^ and via our network of collaborators. Individuals were included based on *SETD1B* variants detected in a research setting or routine clinical diagnostics. Affected individuals were investigated by their referring physicians. Individuals (and/or their legal guardians) recruited in a research setting gave informed consent for their research participation. Those individual research studies received approval from an institutional review board (IRB) (Supplementary Methods). Individuals (or their legal guardians) who were ascertained in diagnostic testing procedures gave informed consent for testing. Permission for inclusion of their anonymized medical data in this cohort, including photographs, was obtained using standard forms at each local site by the responsible referring physicians.

### (Next generation) sequencing of affected individuals

Full details are provided in the Supplementary Methods and Supplementary Figure S1.

### Variant classification

*SETD1B* variants were initially classified as Unknown Significance, Likely Pathogenic or Pathogenic at the performing laboratory or local referring sites. Literature and public database search identified 30 individuals with *SETD1B* sequence variants (Supplementary Table S1). Re-classification of *SETD1B* sequence variants was performed according to ACMG Standards and Guidelines^22^ (Supplementary Table S1), using reference sequence NM_001353345. For retrieval of population allele frequencies and *in silico* predictions the software Alamut® Visual 2.15 (Feb 2020) was used.

### Facial Gestalt and severity scoring analysis

The Face2Gene (FDNA Inc., Boston MA, USA) research application was used using default settings to generate a composite facial gestalt. Details of severity scoring are described in the supplemental methods.

### Structural protein modelling

Sequences were retrieved from Uniprot. SWISS-MODEL^23^ was used to produce homology models, RaptorX^3^ for prediction of secondary structure and disorder, ConSurf^24^ for conservation analysis, and eukaryotic linear motif (ELM^25^) for short linear protein motif assessment. Models were manually inspected, and variants evaluated, using the Pymol program (pymol.org).

### Experimental procedures

For *in vitro* experiments Flag-tagged wild type (a kind gift of David Skalnik, Indiana University^5^) and variant SETD1B protein and HA-tagged ASH2 were overexpressed in HEK293 cells. Protein isolation, Western blotting and immunocytochemistry were performed following standard procedures^26,27^. Proteins for thermal shift assay were expressed in *E.coli* BL21^28^. Genome-wide methylation profiles were obtained as described previously^8^. Further details on experimental procedures and statistical analysis are provided in Supplementary Methods.

## RESULTS

### Molecular spectrum of *SETD1B* sequence variants

A total of 36 individuals with either heterozygous (n=32, n=28 confirmed *de novo*, n=1 inherited from affected parent), compound heterozygous (n=2, bi-allelic inheritance from healthy parents) or homozygous (n=2, siblings, bi-allelic inheritance from healthy parents) *SETD1B* sequence variants were included in this cohort. Thirty-three variants were detected, of which two were recurrent. This includes eight truncating (n=6 nonsense, n=2 frameshift), one extension, one in-frame inversion, and twenty-three missense variants (Figure 1). Nine variants were classified as uncertain significance, ten as likely pathogenic and fourteen as pathogenic. In literature, twenty-six additional (four recurrent) *SETD1B* variants have been reported including seven truncating, one splicing, one extension, three in-frame insertions or deletions, and fourteen missense variants (Figure 1). Variants are distributed along the entire protein (Figure 1), with the majority of (likely) pathogenic missense variants located within the SET domain region.

### Clinical spectrum

The cohort consists of twenty-four males and twelve females, whose age at last evaluation ranged from 1 to 44 years (median 9 years, interquartile range (IQR) 6 - 15 years). Table 1 gives an overview of the core clinical phenotype, and Figure 2 displays the facial appearance of individuals for whom photographs were available. More details can be found in Supplementary Case Reports and Supplementary Figures S2-4.

**Figure 2:** Facial images of affected individuals. Photographs of 16 individuals (plus one affected mother) with indicated *SETD1B* variants. Dysmorphic features included - amongst others - a slightly elongated face, high anterior hair line, thick arched or straight eyebrows, deep set eyes, a prominent nose, and thin upper lips. Lower right corner shows facial composite images for all individuals or only those with a likely pathogenic or pathogenic variant (see also Supplementary Figure S3 and S4) (note: individual 13 and mother were not included in the composite, given the image angle and glasses hindering Face2Gene program analysis).

**Table 1.** Overview phenotypic features <<Table 1.xlsx>>

### Development and neurological findings

Most individuals were born after an uneventful pregnancy at full term, with an unremarkable neonatal period and anthropomorphic measurements in the normal range. Seven individuals (7/31, 23%) had postnatal hypoglycemia. Virtually all individuals (34/36, 94%) showed global developmental delay in early infancy. Notably, individual 14 and 16 without documented developmental delay are the youngest individuals (respectively 2 and 1 years old). Motor development was delayed in thirty-two individuals (32/36, 89%), with independent ambulation acquired between 1.0-4.5 years of age (median 1.6, IQR 1.3-2.5, one individual is non-ambulatory). Motor performance remained an issue, with clumsiness, coordination difficulties, and poor fine motor movements reported. Hypotonia was documented for sixteen individuals (16/35, 46%), often manifesting in the neonatal or childhood period. Language development was delayed in the majority of individuals (33/36, 92%), with first words acquired between 0.5-3.0 years (median 2.0, IQR 1.1-2.1). Five individuals were non-verbal at time of data collection (15%, respectively 2.5, 3, 3.5, 11 and 19 years old), and at least five additional individuals (15%) were able to speak far fewer words than appropriate for their age. Regression of previously acquired skills was reported in nine individuals, especially with regard to language, and was not obviously linked to epileptic activity. At the last investigation, intellectual disability was present in twenty-eight individuals (28/32, 88%), ranging from mild (n=9), to moderate (n=8) and severe (n=4) (not specified n=7). Formal IQ testing was performed in eleven individuals with an average score of 60 (IQR 48-67) (mild). Autistic features were observed in twenty-four individuals (24/36, 67%); other behavioral issues included hyperactivity (13/34, 38%), sleep disturbance (10/32, 31%), anxiety (11/35, 31%), anger or aggressive behavior (11/35, 31%, including self-mutilation for individuals 4 and 10), and obsessive compulsive behavior (individual 7, 26, 30). Epilepsy developed in twenty-eight individuals (28/36, 78%) with a median age of seizure onset of 3 years (IQR 1.0-5.3). Eight individuals remained seizure-free up to an age of 16 years (range 2-16, median 6.0, IQR 4.5-7.3 years). At their onset, the majority of seizures which could be classified were generalized (n=19) and minority focal (n=5), and included motor (n=9) or non-motor (n=13) involvement, with variable development into seizure types over time (Table 1). Seizure frequency varied (sporadic to very frequent) and was at least daily in the majority of patients. Seizures were reported to be fever-sensitive in three individuals. Whereas seizures were (partially) controlled using various antiepileptic drugs in eighteen individuals, seizures responded poorly or remained intractable in seven individuals. Brain MRI (Supplementary Figure S2) was performed in thirty-tree individuals and was often unremarkable (23/31, 74%). Abnormal MRI findings included non-specific minor subcortical white matter hyperintensities (individual 1), cystic encephalomalacia with ventriculomegaly (individual 4), reduced white matter volume and thin corpus callosum (individual 10), bilateral abnormal signals at frontal, temporal, and occipital lobes (individual 16), extensive irregular gyral pattern with reduced sulcation (individual 19), slightly delayed myelination and small heterotopic gray matter (individual 21), periventricular leukomalacia (individual 30, possible due to an underlying hypoplastic left heart disease), and mild diffuse cerebral volume loss with ex vacuo enlargement of lateral and third ventricles (individual 32).

### Additional Findings

Ophthalmological findings included strabismus (n=5), amblyopia (n=2), myopia (n=2), astigmatism (n=3), and cortical vision impairment (n=1). Eight individuals showed gastro-intestinal symptoms, such as reflux, constipation, and feeding problems. Ten individuals had dermatological symptoms, including eczema, rough or dry skin, café au lait spots, and hypo- or hyperpigmentation. A number of individuals displayed skeletal abnormalities, such as scoliosis (n=5), kyphosis (n=2) and joint hypermobility (n=4). (Recurrent) respiratory and urinary tract infections were reported in six individuals. No malignancies were identified. (Truncal) overweight or obesity was present in fifteen individuals.

### Facial appearance

Facial appearance varied from no discernable dysmorphic features (5 individuals) to mild dysmorphic features (31 individuals, 86%) (Figure 2). Dysmorphic features included prominent rounded nasal tip / bulbous nose (n=15), high anterior hairline (n=11), (uplifted) large earlobes (n=10), overfolded superior helices (n=6), low-set ears (n=5), thin upper lip (n=9), pointed/prominent chin (n=6), deep-set eyes (n=5), synophrys (n=4), full cheeks (n=4), elongated/narrow face (n=5) and/or bitemporal narrowing (n=4), and frontal bossing (n=4). Also, tapering fingers (n=5), brachydactyly (n=3), small hands (n=5), and nail hypoplasia (n=4) were reported (Supplementary Figure S3).

### Structural modelling of variants

The eight truncating variants (p.(His8fs), p.(Phe95*), p.(Tyr96*), p.(Glu412fs), p.(Arg1329*), p.(Arg1524*), p.(Gln1666*), p.(Ala1730*)) are likely to be targeted for nonsense-mediated decay, but if not would result in removal of the SET region eliminating catalytic activity. The variants p.(His10Gln) and p.(Glu94Asp) are located in a disordered region preceding the RRM (Figure 1, Figure 3A) and could affect the specificity of the potential interactions mediated by RRM’s N-terminus^29^. The nucleotide inversion leading to p.(Asn113_Asp121delins9) and the substitution p.(Met170Thr) are located in the canonical β_1_α_1_β_2_β_3_α_2_β_4_ RRM region, whereas p.(Gly195Val) is located at the C-terminal loop of α_3_ (Figure 3A). Residues 113-121 are located in the α_1_ helix known to participate in protein-protein interactions in RRM proteins^29^. Furthermore, the RRM domain interacts with the COMPASS component WDR82^5^. Thus, substitution of this 9-residue stretch is expected to severely compromise the RRM fold and its interactions. p.(Met170Thr) and p.(Gly195Val) could affect substrate recognition of RRM because both residues are involved in RNA binding^30^. p.(Thr281Ile) and p.(Thr318Met) are located downstream of the RRM, in a disordered serine, threonine and proline-rich region that contains numerous predicted phosphorylation sites^25^. Hence, p.(Thr281Ile) and p.(Thr318Met) might affect the phosphorylation landscape of this region. Substitutions p.(Arg429Trp), p.(Pro545Arg), p.(Pro698Ser), p.(Pro793Arg), p.(Arg927His), p.(Arg982Gln), p.(Ala1010Val), p.(Ala1129Val), p.(Pro1328Ser) and p.(Arg1424Gln) are all located in the middle region of *SETD1B* predicted to be largely disordered. The middle portions of Setd1 proteins are divergent^1^, suggesting they may have a role in differential genomic targeting of COMPASS through interaction with different targeting proteins. This role might be affected by the mostly non-conservative nature of these substitutions. p.(Ala1129Val), however, is predicted to introduce a non-canonical 5’ splice donor site at nucleotide position c.3384, which would result in a truncated protein p.(Ala1129fs) with the SET catalytic domains eliminated. p.(Arg1748Cys) is located in the WIN motif (Figure 3A) and expected to significantly decrease the interaction between *SETD1B* and WDR5, which is essential for COMPASS assembly and *SETD1B* participation in H3K4 methylation^6^. Substitutions p.(Arg1792Trp), p.(Arg1825Pro), and p.(Lys1827Arg) are located at the interface with the nucleosome (Figure 3A) and therefore likely affect the interaction with histones and the stability of the complex. Variants p.(Ala1901Val), p.(Ala1901Glu), p.(Tyr1941fs), and p.(Glu1948Lys) are located in the catalytic SET domain (Figure 3A). Ala1901 is situated in a loop that is part of the S-Adenosyl methionine (SAM) substrate binding pocket, but is facing away towards an opposing β-strand that is part of the structural core of the SET domain. The substitution of alanine by the larger and negatively charged glutamic acid would create a large stress on the core of the SET domain and potentially disrupt the structural frame maintaining the SAM substrate-binding site and interactions with the adjacent subunits of the complex, whereas alanine to valine substitution introduces a small physicochemical difference which is likely to create some disturbance. The p.(Tyr1941fs) variant would extend the protein, altering the SET domain and post-SET region that are involved in catalysis and cofactor binding thus likely rendering *SETD1B* inactive (Figure 3A). This C-terminal segment is highly conserved^24^. It covers a substantial portion of the binding pocket for histone H3 and the SAM substrate (including the SAM-binding Tyr1943), and three cysteine residues that together with Arg1962 coordinate a zinc atom. Glu1948 is located in a loop adjacent to the histone H3 binding site and, when superimposed to the yeast COMPASS EM structure (PDB ID 6ven) it is found to be close to the DNA binding surface between Set1 and Bre2 (homolog of ASH2) (Figure 3A). The replacement of the glutamic acid by a lysine changes the charge of that side chain and could affect the interactions of this region.

**Figure 3:**
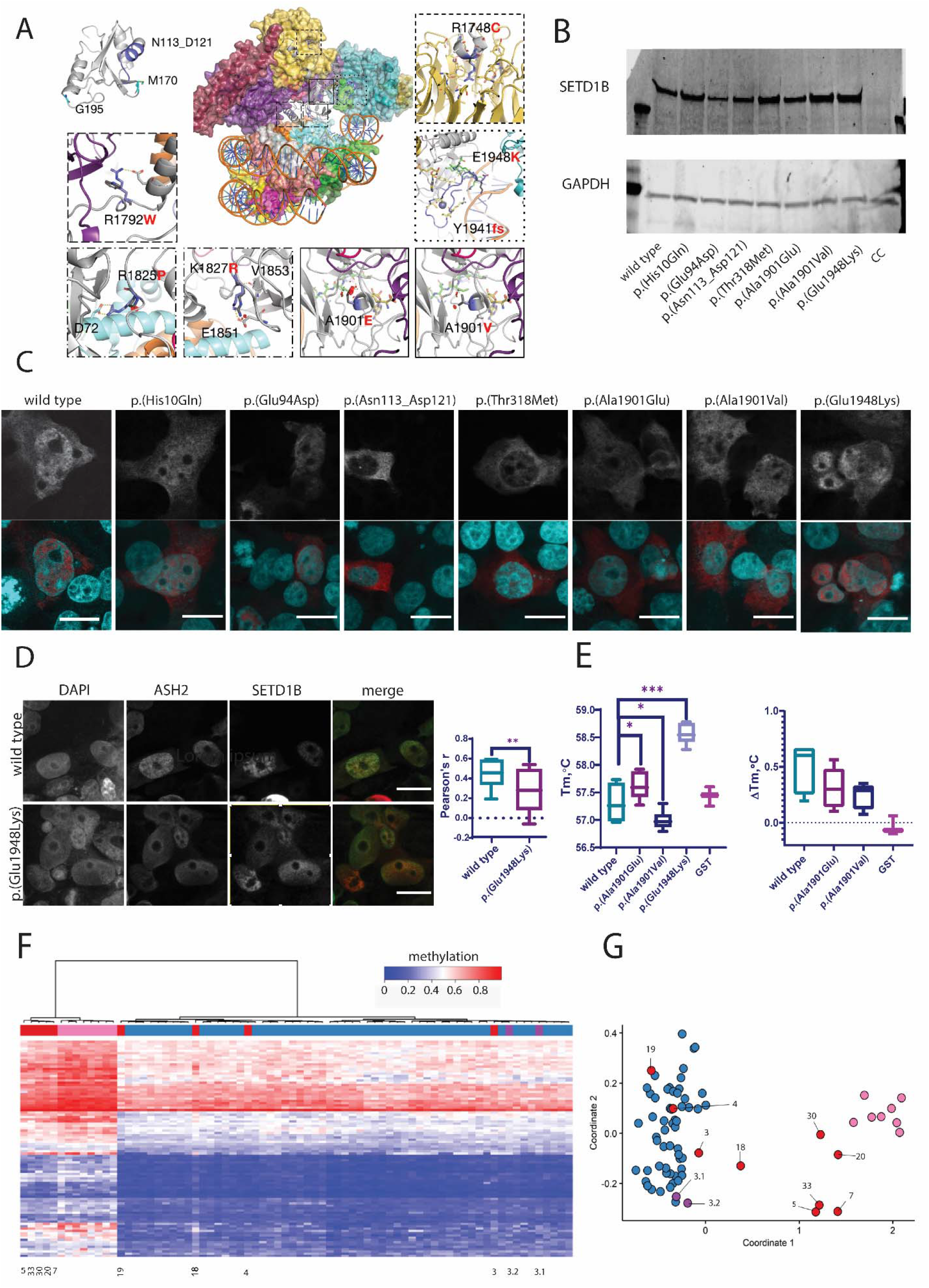
Structural and functional evaluation of SETD1B variants. (A) Homology models of SETD1B domains: RRM domain (top left), based on the crystal structure of the RRM of human SETD1A (PDB ID 3S8S, identity=66%, QMEAN=0.25). The segment of Asn113 to Asp121 is colored in blue. This region is known to support different protein-protein interactions in other RRM proteins. Met170 and Gly195 are shown as blue sticks. Homology model of the N-SET and catalytic SET domains of SETD1B (grey cartoon), based on the EM structure of the yeast COMPASS in a complex with a ubiquitinated nucleosome (PDB ID 6ven, identity= 40.21%, QMEAN=-5.10) (center superimposed to the template PDB, and zoom-in panels). The region containing Arg1748 was observed more accurately in the X-ray structure of the WDR5:SETD1B Win motif peptide binary complex (PDB ID 4es0^6^, top right): Arg1748 (blue sticks) is inserted into the pocket of WDR5 (yellow) and interacts with the backbone oxygen atoms of Ser91, Phe133 and Cys261 (hydrogen bonds shown as yellow dashed lines). Arg1792 (blue sticks) and the substitution by Trp (dark grey sticks) interacting with surrounding residues in the adjacent alpha helix (e.g. Glu1796, grey sticks) or with the SWD1 subunit (RBBP5 in humans, shown in violet). The insets with Arg1825 and Arg1827 show the proximity of these residues (dark blue sticks) to histone H2A (light blue cartoon). The SET domain containing Ala1901Val, Ala1901Glu, Tyr1941fs and Glu1948Lys was modeled more accurately based on the crystal structure of the yeast COMPASS catalytic module (PDB ID 6chg^41^, identity=62%, QMEAN=-1.78). The mutated Ala1901 and Glu1948 are presented as blue sticks in the center figure and right insets. The Ala1901Val and Ala1901Glu substitutions (dark grey sticks) could compromise the stability of the adjacent SAM (olive sticks) binding site and the interaction with the SWD1 subunit (RBBP5 in humans, violet cartoon), which in turn contacts ubiquitin (red cartoon). Tyr1941fs alters a segment of SET and Post-SET regions involved in catalysis and cofactor binding (blue cartoon in center figure and right inset): SAM (olive sticks) and histone H3 (green sticks) binding pocket, the key Tyr1943 residue (yellow sticks), three Cys and one Arg (yellow sticks) coordinating a Zinc atom (shown as a sphere). The Glu1948Lys substitution (blue/dark grey sticks in center figure and right inset) could disturb potential interactions between the flexible loops and the adjacent subunit (Bre2, homologous to human ASH2, is shown in teal cartoon). (B) Overexpression of wild type and variant SETD1B protein in HEK293 cells 48h post-transfection assessed by Western blot. CC-cell control, lysate of mock transfected HEK293 cells. (C) Nuclear localization of SETD1B variants in HEK293 cells. Upper panel – SETD1B detected by anti-Flag antibody; lower panel – overlay of nuclear staining (DAPI, cyan) and SETD1B (red); scale bar 20μm. Images representative of 2 independent experiments are shown. (D) Co-localization of SETD1B and ASH2 in HEK293 cells. Left to right: nuclear staining (DAPI), ASH2 (anti-HA tag), SETD1B (anti-Flag tag), merge of ASH2 (green) and SETD1B (red); scale bar 20μm. Pearson’s r value (range : -1 – negative correlation, 1 – max correlation) calculated with coloc2 plugin (ImageJ), Z-stacks of min. 12 nuclei were used for the analysis. T-test ** p=0.005 (E) Thermal shift analysis of the SET domain. Left – Tm of GST-SETD1B proteins and GST control. Right – change in Tm of the proteins in presence of SAM substrate. Two independent protein preparations were used for the assay performed in triplicates. One-way ANOVA multiple comparison test* p<0.05, *** p< 0.0001 (F,G) Analysis of methylation profiles. – F: Hierarchical clustering (rows represent methylation probes, columns - samples); –G: MDS plot (control samples in blue, proband samples in red, SETD1B cases from the database in pink). Sample numbers correspond to case numbers, 3.1 and 3.2 are the parents of individual 3: p.(Arg982Gln) – case (18), p.(Ala1010Val) - case (19), p.(Glu1948Lys) - case (33), p.(Asn113_asp121delins9) - case (7), p.(Phe95*) – case (5), p.([His10Gln];[Arg927His]) - case (3), p.(Ala1901Glu) - case (30), p.(Ala1129Val) - case (20), p.([Glu94Asp];[Pro1328Ser]) - case (4).

### Functional evaluation of selected SETD1B variants

Based on the structural modeling, seven variants in different regions of SETD1B were selected for *in vitro* studies: p.(His10Gln) and p.(Glu94Asp) N-terminal of RRM, p.(Asn113_Asp121delins9) in RRM, p.(Thr318Met) C-terminal of RRM, and p.(Ala1901Val), p.(Ala1901Glu) and p.(Glu1948Lys) in the catalytic SET domain.

First, stability of *SETD1B* in cells was evaluated by Western blotting of wild type and variant *SETD1B* overexpressed in HEK293 cells (Figure 3B, Supplementary Figure S5A). No significant differences in protein levels were observed, suggesting that the evaluated variants do not affect protein stability (one-way ANOVA p=0.09). Secondly, genomic targeting of *SETD1B* might be dependent on the central region and the catalytic domain, whereas RRM could reinforce chromatin binding,^1,31^ resulting in distribution in the nucleus and not in the nucleoli. Therefore, *SETD1B* nuclear distribution of wild type and variant was assessed by immunofluorescence of transiently transfected HEK293 cells. Overexpressed FLAG-*SETD1B* was detected in the cytoplasm and nucleus. The nuclear localization pattern of *SETD1B* remained similar between wild type and variants, except for p.(Asn113_Asp121delins9) which failed to localize to the nucleus (Figure 3C). Exclusion from the nucleus correlates with an inability to bind chromatin, resulting in loss-of-function of the variant p.(Asn113_Asp121delins9). Thirdly, as suggested by structural modeling based on the yeast COMPASS structure, Glu1948 could be involved in interaction with COMPASS subunit ASH2. Co-transfection and co-localization analysis was performed to evaluate a possible effect of p.(Glu1948Lys) on the respective protein-protein interaction (Figure 3D). Both overexpressed *SETD1B* and ASH2 were detected in the nucleus and cytoplasm of transfected HEK293 cells, with a higher co-localization correlation for wild type compared to p.(Glu1948Lys) variant (Pearson’s correlation value of 0.5 and 0.3 respectively, T-test p=0.005). This evidence for a decreased interaction of p.(Glu1948Lys) with ASH2 provided a rationale for a decreased functionality of this variant. Next, to evaluate the effect of p.(Ala1901Val), p.(Ala1901Glu) and p.(Glu1948Lys), protein stability and ligand binding were evaluated using thermal shift analysis of the catalytic domain (Figure 3E). After GST-tagged *SETD1B* SET-domain expression and purification of wild type and variants, melting temperature (*Tm*) was compared (Figure 3E, left panel; Supplementary Figure S5C, S5D). The *Tm* of p.(Glu1948Lys) was 1.2⁰C higher compared to the wild type (one-way ANOVA p<0.0001), which indicates that this substitution increases the stability of the SET domain which can result in disturbance of interactions within COMPASS, most likely at the interface between *SETD1B*, the nucleosome and the ASH2 subunit, as indicated by the co-localization analysis of this variant with ASH2 subunit (Figure 3D). Substitutions p.(Ala1901Val), p.(Ala1901Glu) resulted in a 0.3⁰C negative and positive shift of *Tm* respectively (one-way ANOVA p<0.05), suggesting that these substitutions have a minor effect on thermal stability and thus on the conformation of the SET domain. However, since these substitutions are predicted to influence the interaction between *SETD1B* and the SAM substrate, the effect on *Tm* in presence of SAM was evaluated (Figure 3E, right panel). Generally, substrate binding stabilizes proteins resulting in an increased *Tm*, and indeed a mean *Tm* increase of 0.3⁰C was observed for wild type. The *T*_m_ changes of the control GST-protein remained <0.1⁰C, suggesting no contribution of GST tag to the SAM interactions. The increase of 0.17⁰C Tm for both p.(Ala1901Val) and p.(Ala1901Glu) indicates no significant effect on SAM interaction of these variants (one-way ANOVA p=0.07 and p=0.19 respectively).

Finally, a specific DNA methylation profile (episignature) for individuals with heterozygous loss-of-function pathogenic *SETD1B* variants has been described^8^. We performed genome-wide methylation analysis for nine individuals (individuals 3, 4, 5, 7, 18, 19, 20, 31, 33), and also the parents of individual 3 with bi-allelic *SETD1B* variants (Figure 3F-G, Supplementary Figure S5F). Individuals 5 (p.(Phe95*)), 7 (p.(Asn113_Asp121delins9)), 20 (p.(Ala1129Val)), 31 (p.(Ala1901Glu), and 33 (p.(Glu1948Lys)) showed the previously established *SETD1B* episignature, individual 18 (p.(Arg982Gln)) showed an inconclusive result, whereas individuals 3 (p.[(His10Gln)];[(Arg927His)]) (nor his parents), 4 (p.[(Glu94Asp)];[(Pro1328Ser)]) and 19 (p.(Ala1010Val)) did not show the *SETD1B* episignature.

Taken together, through structural modeling and functional analyses we provide evidence for reduced function and therefore pathogenicity of p.(Phe95*), p.(Asn113_Asp121delins9), p.(Ala1129Val), p.(Ala1901Glu) and p.(Glu1948Lys), whereas functional consequences and clinical significance remains uncertain for p.(Thr318Met), p.(Arg982Gln), p.(Ala1010Val), p.(Ala1901Val) p.[(His10Gln)];[(Arg927His)] and p.[(Glu94Asp)];[(Pro1328Ser)].

## DISCUSSION

This work reports on the molecular and phenotypic spectrum of 36 individuals with sequence variants in *SETD1B*, representing the largest cohort reported to date. Previous work suggested a possible gain-of-function effect of pathogenic variants in *SETD1B*^14^, however, further reports^8,12,13, 15–19^ including this work, point towards a loss-of-function mechanism. Clinical features of our cohort compared to the eleven previously reported individuals with a (likely) pathogenic *SETD1B* variant^8,12–15^ is provided in Table 2.

**Table 2.**
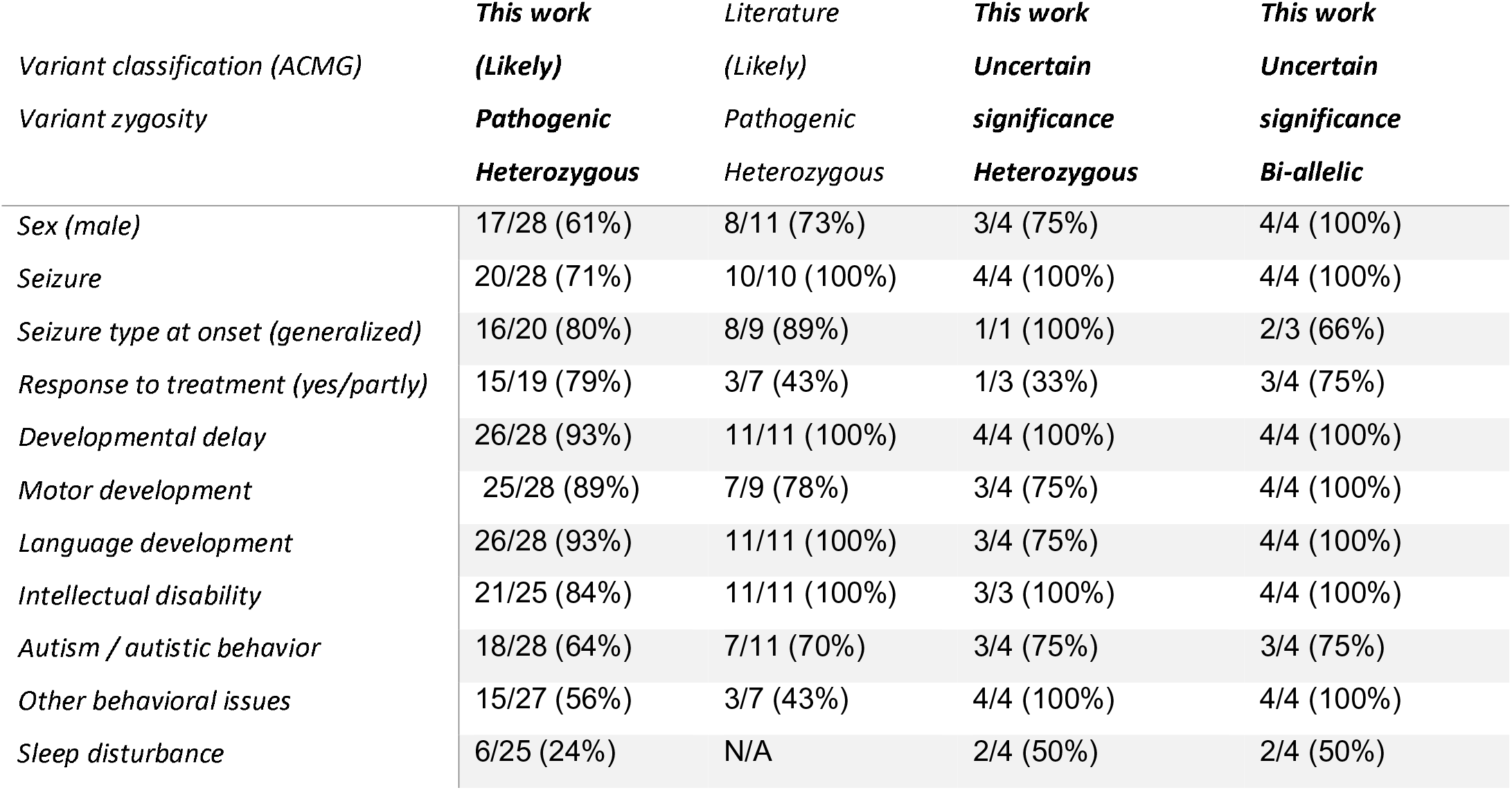
Summary of main phenotypic features in this study cohort and in literature. As information on the different features was not always available for each individual, the denominator of the frequencies differs between the different clinical characteristics.

The emerging phenotype of this *SETD1B*-associated disorder consists of global developmental delay, language delay including regression, intellectual disability, autism, and epilepsy. Other neurobehavioral issues are also often observed, such as hyperactivity, anxiety, anger or aggressive behavior, and sleep disturbance which was not reported before. Importantly, in most cases, developmental delay predates seizure onset, and eight individuals (up to 16 years of age) are without seizures. This indicates that *SETD1B* dysfunction severely impacts physiological neurodevelopment even in the absence of epileptic activity, suggesting the condition is a developmental encephalopathy, with or without epilepsy. Previous work emphasized that alterations of *SETD1B* were mainly associated with myoclonic absences^13^ and predominantly refractory epilepsy. Although myoclonic absence seizures were often observed in our cohort – confirming this association – also other seizure types were regularly encountered at onset, including focal or generalized tonic-clonic seizures. Epilepsy was also well or partially controlled in majority of cases, with just seven (7/26, 27%) remaining refractory to treatment. Brain imaging was unremarkable in most cases, and the observed abnormalities were variable without a consistent phenotype emerging. Whereas previous work did not elucidate consistent dysmorphic features, our cohort identifies a number of mild but consistent dysmorphisms in 30 individuals, including a prominent rounded nasal tip and bulbous nose, high anterior hairline, a thin upper lip, mild ear dysmorphisms, deep set eyes and mild hand abnormalities including tapering fingers, brachydactyly, small hands, and nail hypoplasia. Finally, previous work reported on potential susceptibility to malignancy in *SETD1B*-related disorder^12^, but malignancies were not identified in our cohort, although this remains important for follow-up given the relatively young age of the cohort.

To identify possible genotype-phenotype correlation, a severity score was calculated for each individual in our cohort based on clinical features (Supplementary Methods). No association could be identified between the clinical severity score and the effect or location of the corresponding *SETD1B* variant (Supplementary Figure S6). Intriguingly, there is a significant overrepresentation of males in both our cohort and in literature, with a total of 36 males and 16 females with *SETD1B* sequence variants reported (Binominal test two-tailed p = 0.008, assuming a 50:50 male:female ratio) (Supplementary Methods). The reason for this remains unclear. Incidence of hypotonia and seizures did not differ between males and females in our cohort (hypotonia resp. 12/24, 50% and 4/11, 36%; seizures resp. 19/24, 79% and 9/12, 75%), and seizure onset was similar (resp. range and median years 0-12, 3 and 0-11, 2). Behavioral issues were seen more often in males than in females (autistic behavior respectively 19/24, 79% and 5/12, 42%; hyperactivity respectively 10/23, 43% and 3/11, 27%; anxiety respectively 9/23, 39% and 2/12, 17%; aggression respectively 9/23, 39% and 2/12, 17%; sleep disturbance respectively 8/20, 40% and 2/12, 17%), although differences were not significant between both sexes (Fisher’s test, p>0.05). The combined clinical severity score is significantly lower in females compared to in males (independent sample t-test p=0.025), especially when considering behavioral features as a group (independent sample t-test p=0.006) Supplementary Figure S6). It is thus possible that females present with a milder phenotype which may not prompt medical evaluation. However, ascertainment bias for the neurodevelopmental phenotype could also contribute to the male predominance. Nevertheless, it is tempting to speculate that sex-linked traits could affect susceptibility to clinical penetrance and spectrum of *SETD1B* variants, as female-protective effects have been proposed for other neurodevelopmental disorders^32,33^.

We report four males from three families with bi-allelic variants in *SETD1B*, in which variants were inherited from unaffected parents. Pathogenicity of these variants could not be experimentally proven either by expression, localization or thermostability assays for p.(His10Gln) p.(Glu94Asp) and p.(Thr318Met), nor by methylation analysis for p.[(His10Gln)];[(Arg927His)]) and p.[(Glu94Asp)];[(Pro1328Ser)]. However, this does not exclude the involvement of these variants in yet unknown *SETD1B* functions. Given that the phenotype of these individuals is similar to the heterozygous individuals, (Table 2), and complete absence of *SETD1B* is lethal in several species^10,34,35^, we speculate that the combined action of both alleles in bi-allelic cases results in a phenotype similar to that observed in heterozygous cases by reducing the remaining *SETD1B* activity below a required threshold. A small subset of genes that typically harbor *de novo* variants has already been associated with recessive inheritance^36^. Further investigations remain necessary to establish causality of these variants, and the possibility of recessive inheritance of the *SETD1B*-related disorder.

*SETD1B* adds to a growing list of chromatin modifying genes that have been implicated in neurodevelopmental disorders. *SETD1B* is one of the six H3K4 methyltransferases present in mammals, and remarkably loss-of-function of each is associated with human disease (*KMT2A*: Wiedemann-Steiner syndrome (OMIM #605130); *KMT2B*: early onset dystonia (OMIM #617284); *KMT2C*: Kleefstra syndrome type 2 (OMIM #617768); *KMT2D:* Kabuki syndrome (OMIM #147920)), with the latest additions to this list being *SETD1A* and *SETD1B* (also known as *KMT2F* and *KMT2G*, respectively). *SETD1B* is paralogous to SETD1A (derived from the orthologue Set1) and both associate with the same non-catalytic COMPASS components. *SETD1A* and *SETD1B*, however, show non-overlapping localization within the nucleus and thus likely make non-redundant contributions to the epigenetic control of chromatin structure and gene expression^1^. This might explain why both *SETD1A* and *SETD1B* knock-out mice are embryonically lethal, albeit at different stages of development^34^. Also, in adult mice, *SETD1B* knockout is lethal and was shown to provoke severe defects in hematopoiesis^35^. Heterozygous pathogenic variants in *SETD1A* have been described in individuals with developmental delay, intellectual disability, subtle facial dysmorphisms, and behavioral and psychiatric problems^37^ (OMIM #619056). Interestingly, despite the anticipated non-redundant contributions of *SETD1A* and *SETD1B* to epigenetic control, the clinical phenotype of both related disorders shares many similarities^37^. These include global developmental delay with motor and language delay, intellectual disability, and behavioral abnormalities. *SETD1A* variants have also been found in large scale schizophrenia cohorts^37^ and mouse models support the involvement of *SETD1A* in schizophrenia^38^. One (likely pathogenic) *SETD1B* variant without clinical information was identified in a schizophrenia cohort^39^, but psychosis was not reported in our *SETD1B* cohort. Given the relatively young age of the cohort, this will be an important point for follow-up. Noticeable differences between both syndromes are the incidence of epilepsy which is more common for *SETD1B* (20% in *SETD1A*^37^, 78% in this cohort), the absence of a male predominance for *SETD1A* (9 males out of 19 cases)^37,40^ and differences in the mild dysmorphic features observed in both syndromes (Supplementary Figure S4).

Germline mutants of *Set1,* the orthologue of *SETD1A* and *SETD1B* in *Drosophila melanogaster*, are embryonically lethal^10^, whereas post-mitotic knock-down in neurons shows that *Set1* is required for normal memory in flies, suggesting a role in post development neuronal function^37^. In *Caenorhabditis elegans*, the SETD1A/*SETD1B* orthologue Set-2 is important for transcription of neuronal^9^ genes, proper axon guidance, and neuronal functions, further underscoring the importance of both SETD1A and *SETD1B* for neural function. Interestingly, whereas this work comprises multiple missense variants in the functional domain of *SETD1B* (RRM, N-SET, SET), in SETD1A only one missense variant is reported within a functional domain (post-SET). Finally, of the 23 missense variants found in *SETD1B*, 17 are in regions that are homologous in SETD1A. Of note, p.(Arg982Gln) in the disordered region, is at a homologous position in SETD1A previously described in a patient with early-onset epilepsy (NM_014712.2(SETD1A):c.2737C>T, p.(Arg913Cys))^40^. It will be interesting to decipher the downstream epigenetic alterations causative for the resulting overlaps and differences in phenotype between both syndromes.

## ACKNOWLEDGEMENTS

We would like to thank all patients and families for participation in this study. Part of this research was made possible through access to the data and findings generated by the 100,000 Genomes Project. The 100,000 Genomes Project is managed by Genomics England Limited (a wholly owned company of the Department of Health and Social Care). The 100,000 Genomes Project is funded by the National Institute for Health Research and NHS England. The Wellcome Trust, Cancer Research UK, and the Medical Research Council have also funded research infrastructure. The 100,000 Genomes Project uses data provided by patients and collected by the National Health Service as part of their care and support. Family 2 was collected as part of the SYNaPS Study Group collaboration funded by The Wellcome Trust and strategic award (Synaptopathies) funding (WT093205 MA and WT104033AIA) and research was conducted as part of the Queen Square Genomics group at University College London, supported by the National Institute for Health Research University College London Hospitals Biomedical Research Centre. HH is funded by The MRC (MR/S01165X/1, MR/S005021/1, G0601943), The National Institute for Health Research University College London Hospitals Biomedical Research Centre, Rosetree Trust, Ataxia UK, MSA Trust, Brain Research UK, Sparks GOSH Charity, Muscular Dystrophy UK (MDUK), Muscular Dystrophy Association (MDA USA). GMM was supported by Jordan’s Guardian Angels, the Brotman Baty Institute and the Sunderland Foundation. JRL acknowledges support by the Baylor Hopkins Center for Mendelian Genomics funded by the US National Human Genome Research Institute (UM1 HG006542). The DECODE-EE project (Health Research Call 2018, Tuscany Region) provided research funding to RG. The Epilepsy Society supported this work, with funding to SMS. SMS acknowledges that her work was partly carried out at NIHR University College London Hospitals Biomedical Research Centre, which receives a proportion of funding from the UK Department of Health’s NIHR Biomedical Research Centres funding scheme. A.J. is supported by Solve-RD. The Solve-RD project has received funding from the European Union’s Horizon 2020 research and innovation program under grant agreement No 779257. STA, RR, KJCL, KAPG and FJGV were supported by funding from King Abdullah University of Science and Technology (KAUST) through the baseline fund and the Award No. FCC/1/1976-25 and REI/1/4446-01 from the Office of Sponsored Research (OSR). TSB’s lab is supported by the Netherlands Organisation for Scientific Research (ZonMW Veni, grant 91617021), a NARSAD Young Investigator Grant from the Brain & Behavior Research Foundation, an Erasmus MC Fellowship 2017 and Erasmus MC Human Disease Model Award 2018.

## Author Contributions

Conceptualization: MJAW, TSB

Data curation: MJAW, TSB

Formal analysis: MJAW, KL, TSB

Funding acquisition: TSB

Investigation: functional experiments: KL; methylation analysis: MA, JK, HMC, SH, BS; computational modelling: FJGV, RR, KJCL, KAPG, STA; patient recruitment, clinical and diagnostic evaluations: AJ, YvB, BvP, AK, MvS, NMA, CMK, HR, LP, SB, MZ, SE, HH, IJ, LL, TM, IS, SML, CALR, DQBS, CMPS, HG, NM, SMS, VH, AS, JT, LAP, YLT, IKC, JRL, AGF, JMG, LY, SH, XW, TJ, GR, XL, SM, JLM, GMM, AT, JS, ME, AMP, LS, MRZR, AV, MM, RG, DCK, TMM, MTP, KLMB, JP, ZP, MW, SK, MV, PT, BdV, CG, RP, ML, SRB, KCK, FV, MvG, CQ, MAD, JAH, AMP, RB, LW, SAW, JB, REP, KGM, AC, JKR, AR, HZE, GEL, RM, SB

Methodology: MJAW, KL, TSB

Writing – original draft: MJAW, KL, TSB

Writing – review & editing: all authors

This paper contains supplementary information:

**Supplementary Methods**

**Supplemental Case reports**

**Supplementary Table S1:** Classification of all *SETD1B* variant (cohort and literature)

**Supplementary Table S2:** Oligonucleotides for site directed mutagenesis

**Supplementary Table S3:** Severity scores of *SETD1B* variants in this study according to their clinical phenotype.

**Supplementary Figure S1:** Sanger sequences of selected *SETD1B* variants.

**Supplementary Figure S2:** Brain MRI imaging of selected individuals with *SETD1B* variants.

**Supplementary Figure S3:** Photographs of hands and feet of the indicated individuals.

**Supplementary Figure S4:** Facial gestalt of patients with SETD1A or *SETD1B* variants

**Supplementary Figure S5:** Functional evaluation of *SETD1B* variants

**Supplementary Figure S6:** Average severity score in individuals with heterozygous variants

## Supplementary Methods

### Next generation sequencing analysis and recruitment of individuals

The results from Sanger sequencing confirmation can be found in Supplementary Figure S1, for those nine individuals for which this was available.

#### Individual 1 and 36

Exome sequencing of DNA extracted from leukocytes was carried out for the proband and their parents using Human Core Exome (Twist Bioscience) capture followed by 2*150bp Illumina sequencing. Variant calling was performed with GATK 3.8 and variants were annotated using Alamut-Batch (v1.11). *De novo*, X-linked recessive, homozygous and compound heterozygous variants inherited *in trans* were identified for analysis in the proband using a gene-agnostic trio bioinformatics pipeline. Orthogonal validation was performed by targeted Sanger sequencing of *SETD1B* in all three family members of individual 1.

#### Individual 2 and individual 6

This study was approved by local institutional IRB/ethical review boards (Project ID: 07/N018, REC Ref: 07/Q0512/26), and written informed consent was obtained prior to genetic testing from the families involved. Clinical details were obtained through medical file review and clinical examination.

Genomic DNA was extracted from peripheral blood samples according to standard procedures of phenol chloroform extraction. WES on each proband was performed as described elsewhere^42^ in Macrogen, Korea. Briefly, target enrichment was performed with 2 μg genomic DNA using the SureSelectXT Human All Exon Kit version 6 (Agilent Technologies, Santa Clara, CA, USA) to generate barcoded whole-exome sequencing libraries. Libraries were sequenced on the HiSeqX platform (Illumina, San Diego, CA, USA) with 50x coverage. Quality assessment of the sequence reads was performed by generating QC statistics with FastQC (http://www.bioinformatics.bbsrc.ac.uk/projects/fastqc).

Our bioinformatics filtering strategy included screening for only exonic and donor/acceptor splicing variants. In accordance with the pedigree and phenotype, priority was given to rare variants (<0.01% in public databases, including 1,000 Genomes project, NHLBI Exome Variant Server, Complete Genomics 69, and Exome Aggregation Consortium [ExAC v0.2]) that were fitting a recessive (homozygous or compound heterozygous) or a de novo model and/or variants in genes previously linked to developmental delay, intellectual disability and other neurological disorders.

The family was collected as part of the SYNaPS Study Group collaboration funded by The Wellcome Trust and strategic award (Synaptopathies) funding (WT093205 MA and WT104033AIA). This research was conducted as part of the Queen Square Genomics group at University College London, supported by the National Institute for Health Research University College London Hospitals Biomedical Research Centre.

#### Individual 3

The study was approved by the ethics committees of the Hospital District of Helsinki and Uusimaa and the Institutional review board of Columbia University, New York (IRB-AAAS3433).

For family Individual 3, DNA samples from the affected male individual and both parents underwent exome sequencing. Exomic libraries were prepared using the SureSelect Human All Exon V6 kit (60.46 Mb target region) and paired-end sequencing was performed on a HiSeq2500/4000 instrument (Illumina Inc, San Diego, CA, USA), with an average sequencing depth of on target regions of 68x. Low-quality reads were removed and the filtered reads were aligned to the human reference genome (GRCh37/Hg19) using Burrows-Wheeler Aligner-MEM (BWA)^43^. Duplicate removal, insertions/deletion (Indel)-realignment and base quality score recalibration were performed with Picard-tools and the Genome Analysis Toolkit (GATK). Single nucleotide variants (SNVs) and InDels were called by the GATK HaplotypeCaller^44^. Copy number variants (CNVs) were called in the exome data from the affected individual using CONiFER (v0.2.2)^45^. As part of the quality control, family relations and sex were confirmed using VCFtools and plink^46,47^. SNV/InDel variant annotation and filtering were performed using ANNOVAR^48^ and custom scripts. Variants were filtered by first retaining exonic and splice region variants and based on variant segregation (e.g. autosomal recessive, de novo, X-linked). Next, variants with a predicted effect on protein function or pre-mRNA splicing (missense, frameshift, nonsense, start-loss, splicing, etc.) with a population specific minor allele frequency (MAF) of <0.005 (for AR) and <0.0005 (for AD) in all populations of the Genome Aggregation Database (gnomAD)^49^ were retained. Last, bioinformatic prediction scores were annotated from dbnsfp35a and dbscSNV1.1 to evaluate missense and splice site variants respectively^50,51^. For CNVs, gene annotation was done using the BioMart Database^52^ and variant frequency was assessed using the Database of Genomic Variants^53^ and gnomAD^49^ using the same frequency cut-offs as above for SNV/InDels. SNV/InDel variants were confirmed using Sanger sequencing using an ABI3130XL Genetic Analyzer.

#### Individual 4

Diagnostic trio whole exome sequencing was performed using the AgilentSureSelect v5 capture kit followed by sequencing on an Illumina Hiseq2500 platform (outsourced to GenomeScan, Leiden, The Netherlands). Analysis was performed in the LUMC’s clinical genetic laboratory using an GATK-based pipeline and in-house developed analysis software (LOVDplus).

#### Individuals 5, 7 and 33

Diagnostic trio whole exome sequencing was done as previously described^26^. In short, genomic DNA was isolated from peripheral blood leukocytes of the proband and both parents, and exome-coding DNA was captured with the Agilent SureSelect Clinical Research Exome (CRE) kit (v2). Sequencing was performed on an Illumina HiSeq 4000 with 150-bp paired-end reads. Reads were aligned to hg19 using BWA (BWA-MEM v0.7.13) and variants were called using the GATK haplotype caller (v3.7^44^). Detected variants were annotated, filtered and prioritized using the Bench lab NGS v5.0.2 platform (Agilent technologies). For the Erasmus MC, use of genome-wide investigations in a diagnostic setting was IRB approved (METC-2012-387).

#### Individual 8 and 9

Genomic DNA was isolated from peripheral blood leukocytes. Library preparation was using the TruSeq® DNA PCR-Free Library Prep and sequencing was on the HiSeqX machine. Reads were mapped to GRCh38 using the Isaac aligner and variants were called using the Isaac variant caller, Starling (v2.4.7, Illumina). Variants were then filtered using the Genomics England Tiering process. Individual 8 was recruited in a research study (Protocol number 11/LO/2016 Committee: NRES Committee London – Camden & Islington)

#### Individual 10, 13, 15, 18, 24, 30, 34 and 35

Using genomic DNA from the proband and parents (when available), the exonic regions and flanking splice junctions of the genome were captured using the IDT xGen Exome Research Panel v1.0. Massively parallel (NextGen) sequencing was done on an Illumina system with 100bp or greater paired-end reads. Reads were aligned to human genome build GRCh37/UCSC hg19, and analyzed for sequence variants using a custom-developed analysis tool. Additional sequencing technology and variant interpretation protocol has been previously described^54^. The general assertion criteria for variant classification are publicly available on the GeneDx ClinVar submission page.

#### Individual 11 and 12

Exome capture was performed with the in-house developed BCM-HGSC Core design (52 Mb; Roche NimbleGen, Madison, WI, USA), as previously described^55^. The variant calling was performed by the ATLAS2 suite^56^. Due to suspected consanguinity, the analysis focused on rare homozygous variants shared by the two brothers. Both siblings share five homozygous and two X-linked variants, including variants in three genes previously associated with human disease (*NBAS*, *NOS1*, and *SETD1B*). Bi-allelic variants in *NBAS* are associated with immune defects^57^ and *NOS1* nonsense variants are associated with achalasia^58^. Both variants could hence explain part of the clinical phenotypes of these individuals but not the epilepsy and neurodevelopmental phenotypes.

#### Individual 14, 16 and 25

The exome was captured from peripheral blood DNA using Agilent SureSelectV6 (Agilent Technologies, Santa Clara, California) or IDT xGen Exome Research Panel (Integrated DNA Technologies, Coralville, Iowa). Subsequent paired-end sequencing was using Illumina HiSeq4000 or NovaSeq 6000 (Illumina, Santa Clara, California). Data processing, alignment (using a Burrows-Wheeler algorithm, BWA-mem) and variant calling were performed using Genome Analysis Tool Kit (GATK v4) best practices (https://software.broadinstitute.org/gatk/best-practices/) from the Broad Institute according to the reference gnome GRCh38. Variant annotation was done using ANNOVAR (http://www.openbioinformatics.org/annovar/). Variants in exonic and splicing regions were filter out with a minor allele frequency of ≤0.05 in following databases (1000G, Exome Aggregation Consortium (ExAC), the Exome Variant Server (EVS), the Genome Aggregation Database (gnomAD), and our in-house Chinese population database (CipherDB). Variants were classified as pathogenic (P), likely pathogenic (LP), variant of uncertain significance (VUS), likely benign, or benign in accordance with the guidelines of the American College of Medical Genetics and Genomics (ACMG) and recommendations by The Clinical Genome Resource (ClinGen) Sequence Variant Interpretation (SVI) Working Group (https://www.clinicalgenome.org/).

#### Individual 17, 27 and 29

Saliva samples from patients and their parents were collected (Oragene DNA collection kits, DNA Genotek, Kanata, ON, Canada) and DNA extracted (QIAsymphony, Qiagen, Venlo, Netherlands); blood-derived DNA from the child was also provided by the regional genetics laboratories. DNA samples from patients and their parents were analysed at the Wellcome Trust Sanger Institute with microarray analysis (Agilent 2×1M array CGH [Santa Clara, CA, USA] and Illumina 800K SNP genotyping [San Diego, CA, USA]) to identify copy number variants (CNVs) in the child, and exome sequencing (Agilent SureSelect 55MB Exome Plus with Illumina HiSeq) to investigate single nucleotide variants (SNVs), small insertion-deletions (indels), and CNVs in coding regions of the genome. Putative de novo sequence variants identified using DeNovoGear21 were validated with targeted Sanger sequencing. The population prevalence (minor allele frequency) of each variant in nearly 15[]000 samples from diverse populations was recorded, and the effect of each genomic variant was predicted with the Ensembl Variant Effect Predictor (VEP version 2.6).

#### Individual 19

Paired end reads were mapping to the human genome hg19 using BWA-MEM with default parameters, with reads being additionally processed by The Genome Analysis Toolkit (GATK) and Picard. Variants were identified using haplotype caller within GATK and Freebayes. The intersection of the two variant callers were annotated with SnpEff and loaded into a database using the GEMINI framework. Annotations included predicted functional effect (e.g., splice-site, nonsense, missense), protein position, known clinical associations (OMIM, CLINVAR), mouse phenotypes (MGI), conservation score (PhastCons, GERP), and effects protein function (PolyPhen), CADD scores, and population allele frequencies (Exome Variant Server and Exome Aggregation Consortium data). Tools within GEMINI were used to identify variants confirming to a number of disease models. We focused on variants that are rare in the population (MAF<0.01 or <0.05), are predicted to have a high impact or the gene and are de novo or transmitted in an autosomal recessive, compound heterozygote or x-linked manner^43,44^. (http://broadinstitute.github.io/picard/)

#### Individual 20

Whole-exome capture and sequencing were performed using SeqCap EZ MedExome (Roche NimbleGen). The resulting libraries were sequenced on a HiSeq4000 (Illumina) according to the manufacturer’s recommendations for paired-end 150 bp reads. Alignment of sequence reads to human reference genome (hg19) was done using BWAMEM 0.7.5 (bio-bwa.sourceforge.net/), and variants were called using the GATK3.3 software package (https://gatk.broadinstitute.org/hc/en-us). Filtering of variants was done using Alissa Interpret (Agilent Technologies). Variants with < 5 reads, a frequency of more than 1% in public (ESP, dbSNP, 1KG) and/or in house databases were excluded. *De novo*, homozygous or compound heterozygous variants present in exons or within +/- 6 nt in the intron were evaluated.

#### Individual 21

Trio exome sequencing occurred through Invitae Laboratory (Boosted exome, trio).

#### Individual 22

The study was approved by the Pediatric Ethics Committee of the Tuscany Region, in the context of the DESIRE project (Seventh Framework Programme FP7; grant agreement no. 602531). We performed trio-exome sequencing as previously reported^59^ (Vetro et al, 2020). Briefly, we used the SureSelectXT Clinical Research Exome kit (Agilent Technologies, Santa Clara, CA) for library preparation and target enrichment. We sequenced the captured DNA libraries by a paired-end protocol on Illumina sequencer (NextSeq550, Illumina, San Diego, CA, USA) to obtain an average coverage of above 80x, with 97.6% of target bases covered at least 10x. We performed bioinformatics analysis by standard procedures: we aligned the sequencing reads to the GRCh37/hg19 human genome reference assembly by the BWA software package^43^ and used the GATK suite for base quality score recalibration, realignment of insertion/deletions (InDels), and variant calling, according to GATK Best Practices recommendations^60^. For the annotation and filtering of exonic/splice-site single-nucleotide variants (SNVs) and coding InDels we used commercially available software (VarSeq, Golden Helix, Inc v1.4.6), focusing on non-synonimous/splice site variants with minor allele frequency (MAF) lower than 0.01 in the GnomAD database (http://gnomad.broadinstitute.org/). We further excluded population-specific variants by interrogating our internal database (WES data from over 900 patients with DEE and 200 healthy parents) and evaluated the potential functional impact of SNVs and InDels by the pre-computed genomic variants score from dbNSFP^61^ which was integrated in the annotation pipeline. We also manually interrogated in-silico prediction tools^62–64^, as well as evolutionary conservation scores^65,66^. For selected variants, we visually inspected the quality of reads alignment by using the Integrative Genomics Viewer^67^ and then proceeded to validation by Sanger sequencing (primers and conditions are available upon request).

#### Individual 23

Written informed consent was obtained for all participants in this study under a research protocol approved by the Institutional Review Board at Nationwide Children’s Hospital (IRB18-00662, “Gene Discovery in Clinical Genomic Patients”). Paired-end genome sequencing libraries were constructed for DNA from the proband, mother, and father using NEBNext Ultra II FS DNA Library Prep Kit (New England BioLabs). Whole-genome sequencing was performed on an Illumina NovaSeq6000 instrument according to manufacturer protocols. Reads were mapped to the GRCh37 reference sequence and secondary data analysis was performed using Churchill^68^. The average sequence depth achieved per sample was ∼33x. Our general approach to variant annotation and prioritization has already been described^69^; for this case we prioritized rare nonsynonymous coding variants under several possible inheritance models: Dominant (de novo), recessive (homozygous or compound heterozygous), and X-linked (hemizygous). We identified two de novo coding mutations in the proband: hg19:chr12-122261055-C-T, NM_001353345.2:c.4570C>T *SETD1B*:(p.Arg1524Ter) and hg19:chr1-179314193-T-C, NM_003101.6:c.1099T>C: SOAT1:(p.Phe367Leu).

#### Individual 26 and 28

Diagnostic exome sequencing was done at the Departments of Human Genetics of the Radboud University Medical Center Nijmegen, The Netherlands, and performed essentially as described previously^70^. This study was approved by the institutional review board ‘Commissie Mensgebonden Onderzoek Regio Arnhem-Nijmegen’ under number 2011/188.

#### Individual 31

Trio-exome sequencing of individual 31 occurred as previously described^70^

#### Individual 32

DNA was extracted from peripheral blood using the Promega Maxwell RSC DNA Extraction Kit. The Clinical Exome Sequencing (CES) library was generated using the Agilent SureSelect Human All Exon V6 plus a custom mitochondrial genome capture kit. Captured DNA fragments were then sequenced using the Illumina Nextseq 500 or HiSeq 4000 sequencing system, with 2×100 basepair (bp) paired-end reads. Single nucleotide variants (SNVs) and small insertions and deletions (<10 bp) were detected by mapping and comparing the DNA sequences with the human reference genome (GRCh37/hg19). Variant confirmation by Sanger sequencing is performed for all insertions and deletions as well as substitutions that do not meet the laboratory’s coverage and quality score thresholds.

### Site directed plasmid mutagenesis and in vitro experiments

#### Plasmids

Human full-lengh *SETD1B* construct in pcDNA vector was a kind gift of Dr. David Skalnik, Indiana University^5^. Selected patient variants were introduced by site-directed mutagenesis according to the Q5^®^ Site-Directed Mutagenesis Kit protocol (NEB). To this end the N- and C-terminal parts of *SETD1B* were cloned into pJet vector, mutations were introduced by SDM and then ligated into the original plasmid. All created plasmids were validated by Sanger sequencing. Oligonucleotides used for SDM and sequencing are given in Supplementary Table S2 and plasmid maps are available upon request.

For bacterial protein expression the SET domain of WT or mutant *SETD1B* (amino acids 1727–1966) were cloned into pGEX-4T-1 GST vector (kindly provided by Dr. Mark Nellist, Erasmus MC). All plasmids were verified by Sanger sequencing.

### Overexpression of Flag-*SETD1B* constructs in HEK cells

HEK293 LTV cells were cultured in DMEM medium supplemented with 10% FBS at 37⁰C, 5% CO_2_. Cells were transfected with Flag-*SETD1B* plasmids at ∼70% confluence using Lipofectamine3000 reagent. After 48h cells were processed for analysis by Western blot or fixed in 4% PFA for immunofluorescence staining. Confocal images were acquired with Leica Stellaris5 LIA system using LASX software.

The following antibodies were used: Rb-GAPDH (Cell Signaling, 2118S), anti-Ms-cy2 (JacksonImmunoResearch,711-225-150), anti-Rb-cy5 (JacksonImmunoResearch,711-175-152); Ms-FLAG M2 (Sigma-Aldrich, F3165); Rb-HA tag (Proteintech, 51064-2-AP); IRDye 800CW Goat anti-Rabbit (Li-cor, 926-32211); IRDye 680RD Goat anti-Mouse (Li-cor, 926-68070).

### Bacterial protein expression

pGEX-*SETD1B* constructs were transformed into *E.coli* BL21 competent bacteria. Individual clones were picked and pre-cultures grown overnight at 37°C, 200 RPM. This culture was then used as inoculum to grow bacterial biomass until OD (A600) reached ∼0.7. GST-*SETD1B* expression was induced with 1mM isopropyl β-D-1-thiogalactopyranoside (IPTG) for 20h at 18°C, 200RPM. Cell pellets were resuspended in the lysis buffer 50 mM Tris (pH 7.5), 300 mM NaCl, 10% glycerol, 3 mM DTT, 1 μM ZnCl2), supplemented with cOmpleteTM protease inhibitor tablets (Roche Applied Science)^28^. The cells were sonicated (2 x 30 seconds, 14 µm) and the lysates were cleared by centrifugation. The lysates were incubated at +4⁰C overnight on a rotation wheel with Glutathion-sepharose beads (GE Healthcare). The beads were washed 3x with the lysis buffer and *SETD1B* constructs were eluted in lysis buffer supplemented with 20mM L-glutathione after 10min incubation at RT.

### Thermal shift assay

Thermal shift assay was performed according to a previously published protocol^71^ in a Biorad real-time thermal cycler CFX96 in transparent hard-shell 96-well plates (Biorad). The 25µl reaction contained protein of interest (final concentration 0.2ng/µl), 2.5 µl of 200x SyproOrange dye (Sigma-Aldrich), assay buffer (50mM Tris pH 7.5, 300mM NaCl). For the substrate binding test the reaction contained 100µM of S-(5[]-Adenosyl)-L-methionine chloride dihydrochloride (SAM) (Sigma-Aldrich). Melt curve in the range of 20-90⁰C (increment of 0.2⁰C/10sec, FRET readout) was assessed. The buffer and water controls without proteins were included in the run for background fluorescence.

### Severity scoring

For each individual, a phenotype severity score was calculated by adding up the main phenotypical features from Table 2, which were measured and reported in Table 1. The features taken into account were the following:

*Seizure features*

Seizure: presence of seizure=1, absence=0
*Development features (no=0, any affectation=1)*

- Developmental delay
- Motor development
- Language development
- Intellectual disability (no affectation=0, mild=1, moderate=2, severe=3)
*Behavior features (yes=1, no=0)*

- Autism
- Other behavioral issues (any=1, no=0):

- Hyperactive
- Anxiety
- Aggressive behavior
- Sleep disturbance

The sum was then normalized dividing it by the highest score possible for each individual. The resulting score has a range from 0 (no clinical features observed) to 1 (presentation of all examined clinical features). Scores for only development (see list above) and behavior (see list above) were also calculated by this way.

### Genome-wide methylation profiles and data analysis

Genome-wide methylation profiles were obtained using the Infinium MethylationEPIC BeadChip array (Illumina)^8^. The *SETD1B* EpiSignature has been implemented in the clinical genome-wide DNAmethylation assay, “EpiSign”, and methylation profiles were analysed using the Multiclass Classification Algorithm of EpiSign v2^72^.

Methylation levels calculated as the ratio of methylated signal intensity over the sum of methylated and unmethylated signal intensities, called the β-values, were converted to M-values using logit transformation in order to obtain homoscedasticity for linear regression modeling using the limma package^73^. The model matrix was constructed by these values. The estimated blood cell proportions derived by the algorithm developed by Houseman et al^74^. were added as confounding variables. Subsequently, eBayes function was operated to moderate the created p-values. In order to select the probes, we first selected 1000 probes with the highest product of methylation differences between case and control samples and the negative of the logarithm of multiple-testing corrected p-values derived from the linear modeling by Benjamini-Hochberg (BH) method. Next, a receiver’s operating characteristic (ROC) analysis was performed for every probe and the pairwise Pearson’s correlation coefficient between them was measured. Using the remaining ∼100 probes, hierarchical clustering by Ward’s method on Euclidean distance was performed using the gplots package. Multidimensional scaling (MDS) was done by scaling of the pair-wise Euclidean distances between samples.

### Statistics

Statistical analysis of *in vitro* experiments used one-way ANOVA multiple comparison test or t-test as indicated in the figure legends. Binominal test was used for sex-specific data analysis, using all individuals from this cohort, the mother of Individual 13, and individuals from literature with reported sex^8, 12–15,75^. Fisher’s exact test was used for comparison of other clinical features between male and female patients from this cohort. The severity scores were compared by Welch Two Sample t-test. All statistical analysis was performed in Graph Pad Prism v8 software.

## Supplementary Case Reports

### Individual 1: c.22dup: p.(His8fs)

Individual 1 is a male, born at 38 weeks gestation. Pregnancy was complicated by gestational diabetes and intrauterine growth restriction. There were no perinatal complications. Parents and younger brother are unaffected. Early development was globally delayed. By four years challenging behaviour was prominent (sporadic tantrums, encopresis). He attended mainstream school at age five years with educational support. Cognitive assessment (WISC-IV) at age six-and-a-half years showed a full scale IQ of 57. Currently aged seven-and-a-half years he suffers from intermittent challenging behaviour, and prefers routine but does not meet criteria for autistic spectrum disorder (ASD). His motor skills are progressing e.g. tying laces, ambulating well. Examination revealed dysmorphic features including deep set eyes, short philtrum, dimpled chin, brachycephaly, cupped ear helices bilaterally with large ear lobes and marked tapering of digits. Weight accelerated from birth (2.7kg: 9^th^ centile) to 53.3kg (>99^th^ centile) at seven and a half years (*Figure case 1*). Neurological examination found increased tone in the left ankle. Skin examination was normal.

Seizures began at age six-and-a-half years and consisted of sudden onset, brief (up to 10 second) myoclonic absence seizures, characterised by unresponsiveness with bilateral arm/shoulder jerking and elevation, or sometimes head involvement. Seizure frequency increased over three months, appearing daily (occasionally several per hour). EEG at age 7 years showed brief generalised inter-ictal bursts of spike/polyspike and wave. Photic stimulation elicited a generalised photoparoxysmal response at 11Hz accompanied by a brief absence seizure with accompanying myoclonic activity of the upper limbs and head (*Figure case 1 F*). He was commenced on sodium valproate, which was discontinued due to worsening weight gain. He had two brief generalised tonic-clonic seizures, one of which occurred during valproate wean. Levetiracetam (24mg/kg/day) introduction led to excellent therapeutic response.

MRI brain showed bilateral sub-cortical white matter signal abnormalities (*Figure case 1 D,E*). Chromosomal microarray and Fragile X syndrome investigations were normal. Gene-agnostic trio exome sequencing analysis identified a heterozygous *de novo* novel frameshift variant in *SETD1B* [(NM_001353345.1:c.22dup p.(His8Profs*30)], absent from the Genome Aggregation Database, predicted as likely pathogenic by introduction of a premature termination codon and production of a transcript expected to be degraded by nonsense-mediated decay.

<u>Figure case 1:</u> (**will be made available upon peer review**) A) Front profile: deep set eyes, short philtrum, dimpled chin (B) Side profile: brachycephaly, cupped ear helices and large ear lobes, (C) Tapering of digit with relatively short fifth digit. D) and E), MRI brain showing white mater subcortical hyperintensities bilaterally (noted in 3/9 patients to date). F) Ictal EEG showing brief burst of generalised 3.5-4Hz spike/polyspike and wave [accompanied by a brief myoclonic absence-jerks and brief elevation of the arms (video not available)], which coincided but was not reproduced with photic stimulation. G) Growth chart demonstrating high body mass index prior to AEDs.

### Individual 2: c.22dup: p.(His8fs)

Individual 2 is a currently 10 year old female, born as the 3^rd^ child to non-consanguineous parents from Bangladesh, who presented for genetic analysis at the age of 7 years. Her elder sibling died at the age of 6 years due to an acute abdomen, and family history was positive for a paternal uncle that was diagnose epilepsy of unknown cause which was well controlled. At the first encounter to medical investigations at 4 years of age, she showed delay in motor development, speech delay and impaired cognitive function with regression. Compared to her two siblings, she had always developed more slowly. She developed epilepsy at 4 years of age, that presented with sudden staring and head nodding, lasting for 1-2 seconds and increasing in frequency over time. An initial EEG showed transient bursts of epileptogenic discharges in sleep, but none in the awake state. On a later EEG, runs of high amplitude epileptiform discharges were noted over F3, C3, P3 and Pz. She was initially treated with valproic acid, clorazepate, levetiracetam and risperidon, and is currently treated with valproic acid, levetiracetam and risperidon. Seizures were initially controlled under this treatment regime, but have recently re-appeared at the last investigation at 10 years of age, with an EEG showing generalized and focal discharges. Cognitive functions further regressed and she became emotional labile. At 6 years of age, a single kidney with a neurogenic bladder and a urinary bladder diverticulum were diagnosed. At the last investigation, no dysmorphisms were noted, but she has a wide spread hyperpigmented area involving the right lumbal to upper umbilical region. Her weight was 43 kg and head circumference 53 cm. She presented as playful and happy but sometimes moody. She is able to read and write limited, can talk well but was noted to have has improper emotional behavior and a low intelligence. Whole exome sequencing identified a c.22dup: p.(His8fs) variant in *SETD1B*, which was absent in both parents.

### Individual 3: c.30C>A, p.(His10Gln); c.2780G>A, p.(Arg927His)

Individual 3 is a 21 years old male, the only child of the family. He was born by urgent caesarean section due placental ablation. He was diagnosed to have asphyxia, pH 6.86, from which he recovered well. His birth weight was 2930 gram, with length of 49 cm and head circumference of 35.5 cm. As a newborn he was studied due slow growth (height -4 SD, weight -1 SD). His developmental delay was noticed at 1.5 years of age. Focal epilepsy started at 2 years of age which developed to Lennox-Gastaut syndrome at 3 years of age for which Absenor and Lamictal were prescribed. He learnt to walk at 3 years of age. He speaks some words. Severe intellectual disability was diagnosed at 4 years of age. His adult height is 158 cm. He has difficulties in walking due hypotonia. His balance is weak and he has a tendency to fall and hurt himself. He has also intention tremor. At 11 years of age aseptic coxitis was detected. His facial features include a narrow skull, narrow and high palate and small, low set ears. His fingers are short and he has a sandal gap between 1. and 2. toes. He had enuresis until 18 years of age. His kidney ultrasound is normal. In EEG interictal spikes are characteristic (Figures Case 3). When awake occasional spikes from both hemispheres, especially from centroparietal areas are detected, whereas during sleep abundant multifocal spikes, especially from centroparietal areas (right side often more affected) are found. In EEG background no reaction to eye opening or sleep phenomena were found (Figure Case 3). Findings were similar in repeated recordings over the years. Brain MRI and ENMG were normal. Screening of urine amino-acids, oligosaccharides and glycosaminoglycans was normal. A HumanCytoSNP-12 (v2.1) (Illumina) was normal. Subsequent trio exome sequencing identified compound heterozygote variants (p.His10Gln and p.Arg927His) in the *SETD1B* gene. The parents are healthy carriers of these variants.

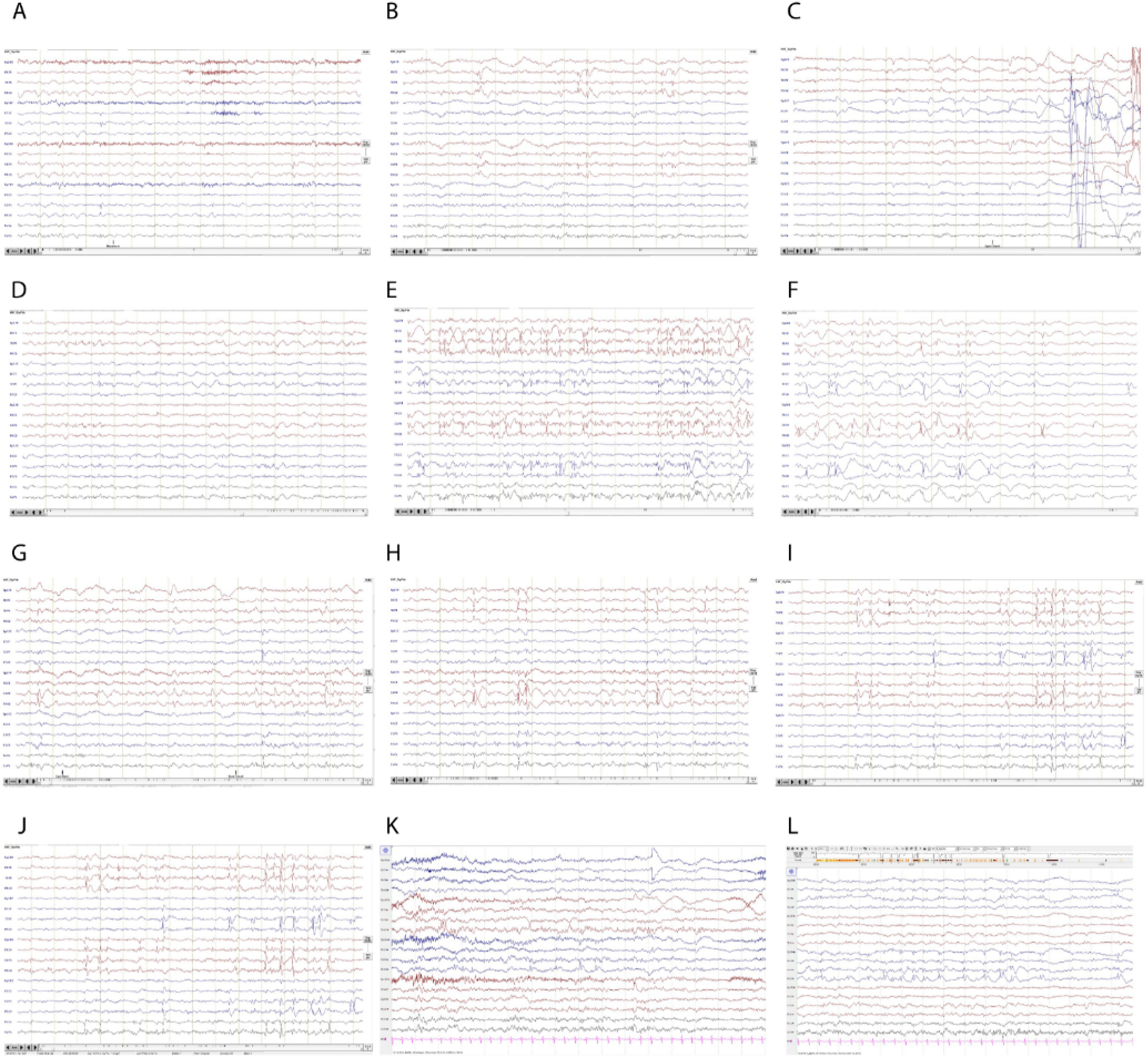

<u>Figure Case 3:</u> Repeated EEG recordings over the years, at awake and sleeping state, showing mainly multifocal spikes at the centroparietal areas. A) and B) age of 6 years, awake, showing occasional multifocal spikes especially at the centroparietal areas. C) age of 7 years, showing continuous EEG background, without posterior rhythms during eye closure. D) age of 6 years, possible during sleep, showing spikes at the centrotemporal areas. E) and F) during sleep at the age of 6 years, showing abundant spikes from the centroparietal areas. G) Continuous EEG at the age of 8 years, awake; H) age 8 years, possible sleeping. I) and J) 9 years of age during sleep. K) age of 10 years, awake and L) age of 10 years, during sleep, showing right parieto frontal spikes (P4-F4), and left sides multifocal spikes (P3, C3, T5).

### Individual 4: c.282G>C, p.(Glu94Asp); c.3982C>T, p.(Pro1328Ser)

After an uneventful pregnancy and home delivery, a boy was born at gestational age of 41+1 weeks with a birth weight of 4.5 kg. On day 1 after birth he experienced episodes of apnea with desaturation and tonic seizures. A cerebral ultrasound and MRI were performed, showing cystic encephalomalacia at the right side of the brain with bilateral ventriculomegaly and displacement of the brainstem. Fenestration of the cyst occurred and an Ommaya shunt was inserted. Seizures were treated with phenobarbitone for 2 months. At the age of 11 months focal motor seizures reoccurred, and antiepileptic drugs were restarted (valproic acid and clobazam). In the year thereafter, valproic acid was switched to oxcarbazepine with good effect. Because of exotropia, he had strabismus surgery at the age of 6 years. He developed some form of speech (copying words and singing short lines) from the age of 2 years, but lost this ability once he started walking. His motor development was severely delayed (ambulation at 4 years). Currently, at the age of 11, he produces sounds but no words. Furthermore, he has pronounced joint hypermobility and severe constipation, for which a high dose macrogol is required. He has autistiform behavior but does not meet diagnostic criteria. Behavioral and sleeping problems seem to increase throughout the winter period.

### Individual 13: c.1234del, p.(Glu412fs)

Individual 13 is a 5 year old male with a dual diagnoses of *PTEN*-related disorder and *SETD1B*-related disorder. He is also a carrier of sickle cell trait. His associated features include macrocephaly, rapid growth, developmental delay with regression, epilepsy, and severe autism.

Whole exome sequencing, which revealed a reclassification of the *PTEN* variant from variant of uncertain significance to likely pathogenic, as well as a maternally inherited pathogenic variant in *SETD1B* <u>(c.1234del, p.(Glu412fs)).</u> It also confirmed his previously known sickle cell trait. The mother is similarly affected with borderline intellectual functioning (IQ72), seizures, and autism spectrum disorder; she is immature for age and has prominent facial features and some slight facial asymmetry, and shares her son’s very straight eyebrows. She is not unaffected, though clearly more high functioning than her affected son. The maternal grandmother did not carry the *SETD1B* variant, but reports a history of learning disability and childhood seizures in herself. She currently functions as the guardian of daughter and grandson.

Of note, the patient is also status post normal chromosome microarray, fragile X testing, MPS screening, inborn errors of metabolism screening, and brain MRI. Most recent thyroid ultrasound was normal.

### Individual 14: c.1285C>T, p.(Arg429Trp)

Individual 14 is a currently 2 years and 3 months old female, born to non-consanguineous Chinese parents, at 40+2 weeks of gestation, with a birth weight of 2.7 kg and a length of 50 cm. Developmental milestones were achieved normally, and no dysmorphic features were noticed. A first seizure occurred at the age of 1 year and 11 months old. Her symptoms started with vomiting and diarrhea, followed by her eyes turned upwards, lips and face turned to pale, generalized convulsions lasted for one minute. Second seizure occurred again recently, with similar symptoms: eyes turn up, left upper limb flexed and raised, chewing, salivation, cyanosis of the lips, and then the whole body twitches, which lasted around 90 seconds. She was treated with phenobarbital orally, and is seizure free so far. Brain imaging was unremarkable.

### Individual 15: c.1634C>G, p.(Pro545Arg)

Individual 15 is a 19 year-old male presented to the Genetics Clinic at 18 years of age, referred by his neurologist for consultation due to his diagnoses of autism spectrum disorder, attention deficit hyperactivity disorder, seizure disorder, and mild intellectual disability. Prior to his Genetics visit, his neurologist ordered a karyotype that revealed an apparently balanced reciprocal translocation between chromosomes 12 and 19 - 46,XY,t(12;19)(p11.2;p13.1). He had a normal microarray. The patient was adopted at 6 months of age from Russia and there is no prenatal history or early developmental history known prior to 6 months. He had onset of seizures at age of 6 years and currently has absence seizures occurring around once a month (well-managed on Keppra and Depakote). He is generally non-dysmorphic on physical exam. Fragile X testing and an epilepsy panel were ordered following his Genetics visit and were both negative. At that time, whole exome sequencing was ordered and a variant of uncertain significance (c.1634C>G, p.(Pro545Arg) was identified in the *SETD1B* gene.

### Individual 16: c.2092C>T, p.(Pro698Ser)

Individual 16 is a currently 1 year old female, born as the first child to non-consanguineous Chinese parents at 38+1 weeks of gestation, with a birth weight of 2.26 kg and a good start (APGAR 10/10/10). Delivery was by cesarean section due to mother’s preeclampsia. After birth, she was hospitalized for treatment because of neonatal pneumonia, hyperglycemia, conjunctivitis, low birth weight and patent arterial duct. One day before she was admitted to hospital at the age of 3 months and 12 days, she had high fever (39.6 C) with unknown cause. Parents tried to physically lower her body temperature by rubbing with alcohol and treated her with antibiotics, and her body temperature decreased to normal gradually. 12 hours before admission, she started with sighing breath, lasted for 5-10 minutes, followed by seizures unconsciously with right upper limb convulsion, left upper limb and both lower limbs tonic-clonic seizures, while head moving leftwards, both eyes gazing leftwards, with a pale face, purple lips, and tightly closed mouth. Seizure lasted about 10 minutes, and her body temperature was 37.9 C at that time. She was admitted to the hospital unconsciously. Lung CT showed signs of a bi-lobular pneumonia and blood work suggested mild anemia and septicemia. She can raise her head steadily by 3 months, is able to smile, but not very active, not very good at following light or moving objects. Her physical examination showed hypertonia, with high deep tendon reflexes (knee and Achilles) and a possibly positive Babinski signs. No dysmorphic features were observed. Brain MRI showed bilateral abnormal signals at temporal, occipital lobes. Current development at 1 year old seems normal. Whole exome sequencing identified a *de novo* c.2092C>T, p.(Pro698Ser) variant in *SETD1B*.

### Individual 21: c.3985C>T, p.(Arg1329*)

Individual 21 is a 4-year-old female with predominantly myoclonic epilepsy and developmental delays. Her early development involved slight delays in walking and language, and typical fine motor skills. Speech and language assessments have shown slightly below average expressive and receptive language skills. She receives occupational and speech therapy. She has not demonstrated any developmental regression. The subject was initially diagnosed with absence seizures with eyelid myoclonia at 2 years old, which evolved into very frequent treatment refractory epilepsy. She typically has around 70-100 myoclonic seizures per day; these frequent myoclonic seizures have been demonstrated on EEG. Prior AEDs trialed include Lamicatal, Diamox, which seemed to possibly increase seizures, and Onfi. She is currently treated with Keppra and Epidiolex, and is on a modified Atkins diet. She seems to have more seizures in the morning or evening and during sedentary activities, like watching tv. Brain MRI at 4 years old revealed right choroid fissure cyst, and subtle features of increased FLAIR signal that may represent slightly delayed myelination and questionable small heterotopic gray matter. The child has not yet had a full dysmorphology exam in clinic. Over telehealth appointment she was noted to have light blond hair, normally shared ears, normal palmar and foot markings, typical gait, and typical facies. Her prior genetic work-up included an epilepsy panel with a variant of unknown significance in the gene CHRNB2, which was considered non-diagnostic. She was also identified to be a Fragile X premutation carrier (57 and 29 CGG repeats). Trio exome sequencing identified a *de novo c.3985C>T, p.(Arg1329*)* nonsense variant in *SETD1B*.

### Individual 22: c.4271G>A, p.(Arg1424Gln)

Individual 22 is a 3 years and 6 months old boy, first child of healthy non-consanguineous Italian parents. Pregnancy was unremarkable and delivery was by induced labor at 39+6 weeks. Birth weight (3615 gr, +0.11 SD), length (51 cm, +0.32 SD) and head circumference (35 cm, -0.40 SD) were within normal range. Global hypotonia and developmental delay were noticed since age of 4 months. The child achieved head control at 1 year and 3 months and independent walking at 3 years. Neuropsychological evaluation at 2 years and 4 months was consistent with moderate developmental delay (Bayley III scale: Cognitive, 55; Language, 49; Motor, 42). Two brain MRI scans, performed at 1 year and 3 years, were normal. At age 3 years, the boy was still non-verbal and manifested brief episodes of unresponsiveness with eyelid myoclonia, accompanied by generalized spike-wave discharges. Treatment with valproate was rapidly beneficial.

Plasma metabolic workup and SNP-array analysis were normal. Whole exome sequencing uncovered the *de novo* c.4271G>A [p.(Arg1424Gln)] *SETD1B* variant of (accession nr. NM_001353345.1), which was classified as likely pathogenic according to ACMG (PS2, PM2, PP3). The variant was not observed in the allele frequency database GnomAD (v2.1) and was predicted to be damaging by multiple *in silico* bioinformatics tools (SIFT, Polyphen2, MutationTaster).

### Individual 23: c.4570C>T, p.(Arg1524*)

Individual 23 is a currently a 5-year-old Caucasian male born at 38 weeks gestation. The pregnancy was complicated by maternal factor V Leiden deficiency, for which mother took baby aspirin during the pregnancy. Other maternal medications included progesterone injections until 12 weeks. Intrauterine growth retardation was noticed, and labor was induced at 38 weeks due to this. Patient weighed 5 pounds, 7 ounces at birth (2.466 kg, 4.19%, Z=-1.73), and was 17.5 inches long (44.5 cm, 2.0%, Z=-2.05). Global developmental delays were noted early on, with the patient not sitting up on his own until 9 months, or crawling until a year of age. Patient began receiving therapies (physical and occupational) at 9 months of age. More severe delays in speech were noted as patient got older, and speech therapy was engaged. Patient began speaking around 2 years of age, and was still using only single-words at 3 years of age. He was diagnosed with apraxia, global developmental delays, and ADHD. At age 4, he had a full developmental and psychological assessment confirming his diagnoses of global developmental delay and ADHD. He had a full-scale IQ of 63 on the Leiter-3 scale at that time. Medically, he has been generally healthy, but did have conductive hearing loss due to fluid in his ears, for which he received ear tubes, and enlarged tonsils, for which he had a tonsillectomy. His current growth parameters at age 5 years, 10 months are 53 pounds, 5.6 ounces (24.3 kg, 86.86%, Z=1.12) and 47.72 inches tall (121 cm, 90.35%, Z=1.30).

The patient, along with his sister and parents, were enrolled in a research study for whole genome sequencing, and the c.4570C>T, p.(Arg1524*) variant in *SETD1B* was found to be *de novo* in the male patient. Parents are both healthy with normal cognition, and there is no additional family history of intellectual disability or birth defects. Consanguinity was denied.

### Individual 24: c.4996C>T, p.(Gln1666*)

Individual 24 is a 14-year-old male with generalized epilepsy and mild intellectual disability. His early development was notable for expressive language delay; gross motor and fine motor skills were attained towards the end of the normal range. Notable challenges in academic skills became apparent with initiation of formal schooling, primarily affecting writing, comprehension, and memory recall. There has been no history of developmental regression. At 6 years of age, it was recommended that he undergo EEG, which demonstrated brief absence seizures occurring every 30-90 seconds. Given their subtle semiology, seizure onset is not known but is thought to be prior to 6 years. The child has since trialed multiple antiepileptic drugs (AEDs) and had a vagus nerve stimulator (VNS) placed at 11 years of age. He continues to have daily clusters of seizures upon awakening with his current AED regimen of cannabidiol, clobazam, and ethosuximide. Known seizure triggers include dehydration, sleep deprivation, and emesis. The subject continues to have difficulty with schoolwork and his most recent neuropsychiatric evaluation revealed a full scale IQ consistent with mild intellectual disability. His neurologic exam was also significant for hypotonia (appendicular > axial) and intermittent tremor. Several minor anomalies were noted on dysmorphology exam, including: low anterior hairline, bilateral posterior helical ear pits, mild synophrys with broad eyebrows, upslanting palpebral fissures, an upturned nose with broad nasal tip, and wide mouth. His prior genetic work-up included a SNP chromosomal microarray and a large epilepsy gene panel. Trio whole exome sequencing identified a *de novo* c.4996C>T, p.(Gln1666*) variant in *SETD1B*.

### Individual 27: c.5374C>T, p.(Arg1792Trp)

Individual 27 is a currently 13 year old female, born at full term weighting 3033 gram, to non-consanguineous parents. During the first 4-6 post-conception weeks of pregnancy mother took recreational drugs (cocaine, cannabis, ecstasy) and alcohol but stopped immediately on learning she was pregnant. Parturition was prolonged and she was a vaginal birth with occiput-posterior. Placenta accreta was noted but there were no neonatal problems and she was breast-fed for 12 months. She was a placid infant who fed well, sat independently at 9 months, walked independently at 20 months, and was late developing babble and speech. She was noted to be hypotonic in infancy and through childhood, with longstanding coordination difficulties and mild ataxia. Her seizure disorder probably started around 6 months of age with absences and was formally diagnosed at 18 months. While presenting mainly as absences her generalized seizure disorder (as suggested by her EEG) can include tonic-clonic episodes. The absences can be very frequent and have been difficult to control, with no significant improvement on either sodium valproate, lamotrigine, or levetiracetam. Some improvement has been manifest on ethosuximide and topiramate. She has mild intellectual disability and speech delay, and communication and social difficulties thought to be consistent with autistic spectrum disorder; however, on formal assessment she was considered not to have autism. She manifests hyperacusis, and has outbursts of hyperactivity as well as aggression. She is medicated for constipation and has episodes of fecal soiling. She has always fed well and been well grown, with height and weight approximately 95^th^ centile throughout childhood. Head circumference was 51.0 cm at age 6 years and 2 months. Parental heights differ markedly: mother ∼138 cm (OFC 55.8 cm); father ∼188cm (OFC 55.7cm). She is not obviously dysmorphic but has mildly anteverted nares and slightly short fingernails and terminal phalanges. She was noted to have mild 5^th^ finger brachyclinodactyly when younger, and at age of 6 years her palpebral fissure length was on the 5^th^ centile. Through the Deciphering Developmental Disorders project she was found to have the *de novo* heterozygous missense *SETD1B* variant: c.5374C>T, p.(Arg1792Trp).

### Individual 30: c.5702C>T, p.(Ala1901Val)

Individual 30 was a male born at 40 2/7 weeks gestation to a 31-year-old female. A fertility herbal blend was ingested by the mother and the father prior to conception. Medications during pregnancy included prenatal vitamins, Valtrex, Glyburide, and Zofran. Hypoplastic left heart syndrome was identified at 22 weeks gestation by fetal echocardiogram. Pregnancy was also complicated by maternal gestational diabetes.

He was born by induced vaginal delivery with a birth weight of 3340 gram, length of 51.5 cm, and head circumference of 33.5 cm. On a postnatal echocardiogram, the diagnosis of hypoplastic left heart syndrome was confirmed, in addition to identification of a moderate-sized atrial septal defect, severe hypoplasia of the ascending aorta (4 mm), moderate hypoplasia of the aortic arch, and patent ductus arteriosus. On day 6 of life, he underwent a hybrid procedure with bilateral pulmonary artery banding and placement of a single ductal stent. At 3.5-months-old, he underwent a multi-procedure surgery that included tricuspid valve repair, removal of ductal stent, patch enlargement, atrial septectomy, Damus-Kaye-Stansel connection, right-sided bidirectional Glenn shunt, and balloon dilatation of the left pulmonary artery. Completion Fontan procedure was performed at 2-years-old. Additional cardiac catheterizations were performed at 1-month-old, 3-months-old, 2-years-old, and 3-years-old. He was hospitalized once after the completion Fontan procedure due to an upper respiratory infection. His cardiovascular status has remained stable.

He experienced developmental delay. He rolled over at 6 months, sat by himself and crawled at 12 months, walked unassisted at 18.5 months, and spoke his first word at 18 months. Verbal regression characterized by the loss of approximately 30 words occurred around 24-months-old. He was diagnosed with autism spectrum disorder by a developmental pediatrician at 3-years-old. Brain magnetic resonance imaging identified mild lateral and third ventricular enlargement with mild periventricular white matter thinning and minimal periventricular white matter FLAIR hyper-intensity suggesting a mild form of periventricular leucomalacia, in addition to several foci of susceptibility artifact scattered in both cerebral hemispheres.

Family history was significant for early-onset breast cancer. His mother was diagnosed with unilateral breast cancer at 28-years-old. Family history was negative for additional cases of congenital heart defect, developmental delay, and autism spectrum disorder. There was no reported consanguinity or Ashkenazi Jewish ancestry.

He had a negative karyotype, microarray, Fragile X testing, and FISH for 22q11.2 deletion. Clinical exome sequencing reported a de novo novel missense variant, c.5702C>T: p.Ala1901Val, in *SETD1B*.

### Individual 32: c.5820_5826del, p.(Tyr1941fs)

Individual 32 is a currently 15-year-old boy recently diagnosed with a *de novo SETD1B* c.5820_5826delCTATGAC, p.Tyr1941IlefsTer101 variant. He presented at 1-1/2 years of age with medically intractable Lennox-Gastaut syndrome. He was noted to be globally developmental delayed since infancy and is affected with intellectual impairment and autistic features. Currently he is becoming less verbal and there is a concern for a regression of expressive language skills. He has fuller cheeks, tapered fingers, thoracolumbar scoliosis and pes planus.

### Individual 34: c.5842G>A, p.(Glu1948Lys)

Individual 34 presented to the hospital as 18 days old male with apneic episodes associated with micrognathia, laryngomalacia, and symptoms of gastoesophageal reflux disease. Laryngomalacia was corrected with supraglottoplasty at 3 months old. By 23 months old, he demonstrated motor and language delay and had been diagnosed with sensory processing disorder. MRI of the brain performed due to history of suspected vestibular dysfunction demonstrated normal results, and there were no abnormalities on an EEG that was conducted during a sleep study. At 30 months old the patient had been formally diagnosed with autism spectrum disorder and did not have history of seizure activity.

### Individual 35: c.5842G>A, p.(Glu1948Lys)

Individual 35 has a history of early developmental delay, medically refractory absence epilepsy with a myoclonic component, and autism spectrum disorder. He was delivered at term with a birth weight of 7 pounds 11 ounces. He experienced early GE-reflux and underwent a frenectomy as an infant. He had early developmental issues and did not walk until starting early intervention therapies at 14 months of age. He also had a significant language delay. His parents began to notice brief staring spells sometimes with a larger jerk in his infancy. This was subsequently diagnosed as epilepsy around the age of 3 years. He was found on EEG to have generalized polyspike spike-wave discharges. Since then he has had medically refractory absence epilepsy with a myoclonic component but also has had generalized tonic-clonic seizures. He was thought initially not to have autism, but as he got older his processing issues and perseverative behaviors became more evident and he was diagnosed with autism spectrum disorder at the age of 4. He has never had regression.

## Supplementary Figures

**Supplementary Figure S1:**
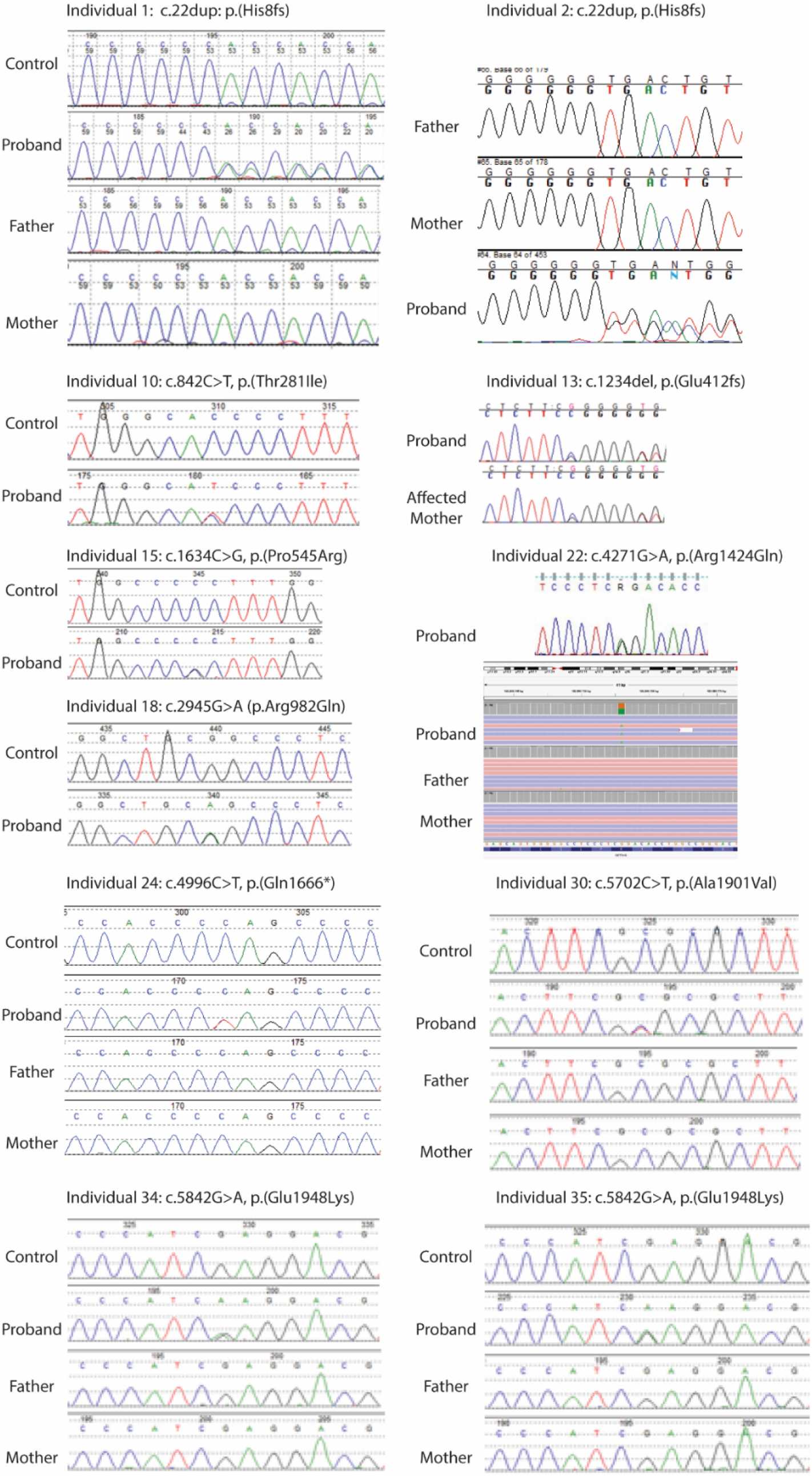
Sanger sequences of selected *SETD1B* variants. Shown are the chromatograms of affected individuals and family members, as indicated. For individual 22, also the IGV view of the trio is shown.

***Figure will be available upon peer review***

**Supplementary Figure S2:** Brain MRI imaging of selected individuals with *SETD1B* variants.

Shown are representative MRI brain images (T1 or T2 weighted) in different planes, for individuals 1, 4, 10, 13, 16, 18, 22 and 24, showing non-specific minor subcortical white matter hyperintensities (individual 1), a cystic encephalomalacia in right hemisphere with ventriculomegaly and the need for a shunt (individual 4), reduced white matter and thin corpus callosum (individual 10), bilateral abnormal signals at temporal and occipital lobes (individual 16), or unremarkable findings (individual 13, 18, 22, 24).

***Figure will be available upon peer review***

**Supplementary Figure S3:** Photographs of hands and feet of the indicated individuals.

***Figure will be available upon peer review***

**Supplementary Figure S4:** Facial gestalt of patients with *SETD1A* or *SETD1B* variants

A. Composite Facial gestalt (as in Figure 1), in comparison to individuals with variants in *SETD1A*. The composite image for *SETD1A* was generated by analyzing all previously published images from Kummeling et al^37^.
B. Comparison between *SETD1A* individuals and individuals with likely pathogenic or pathogenic variants in *SETD1B*.
C. Comparison between *SETD1A* individuals and individuals with likely pathogenic, pathogenic or variants of unknown significance in *SETD1B*.

**Supplementary Figure S5:**
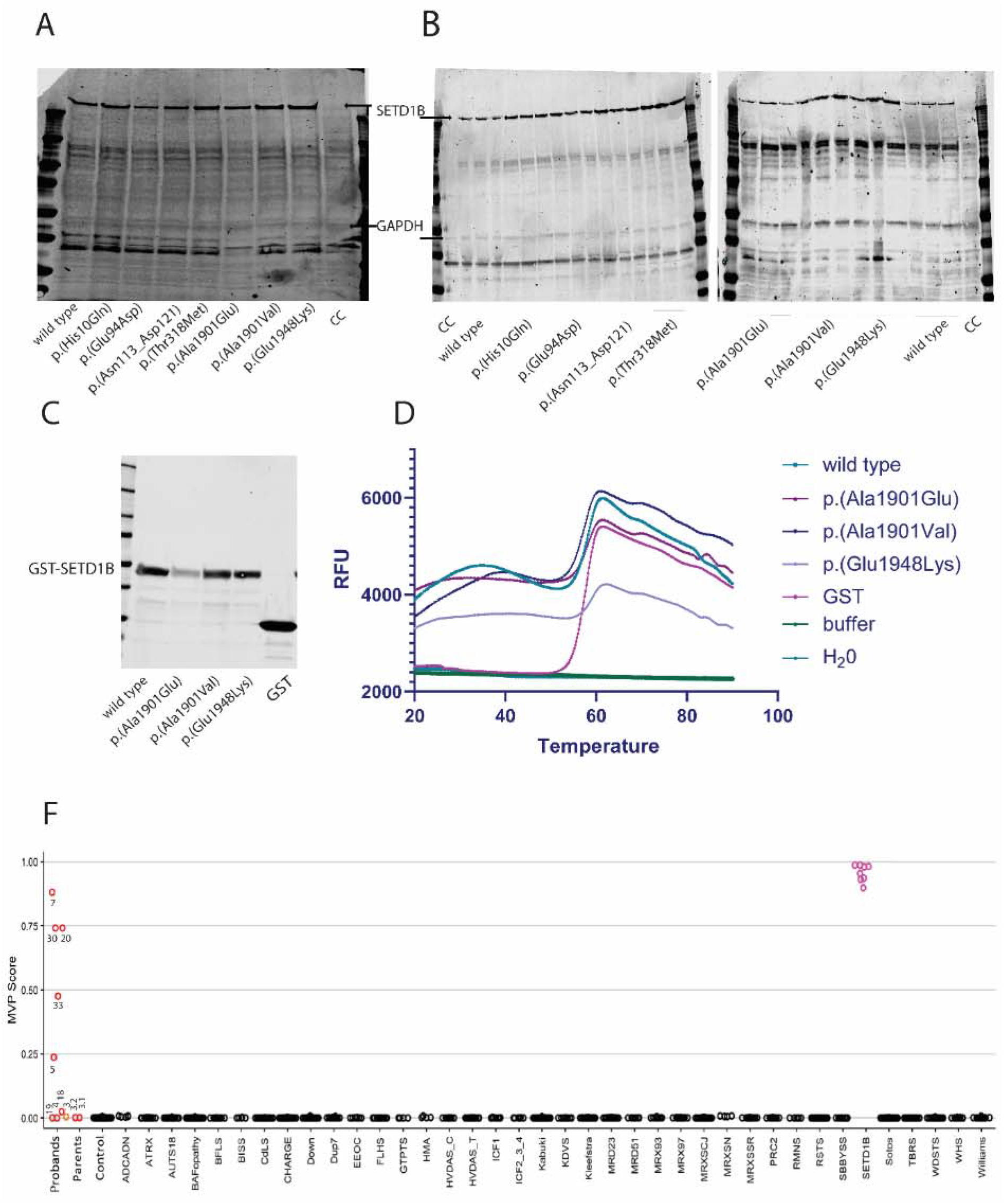
Functional evaluation of SETD1B variants. (A) Full western blot (triplicate) of overexpression of wild type and variant *SETD1B* protein in HEK293 cells 48h post-transfection assessed by Western blot. CC-cell control, lysate of mock transfected HEK293 cells. (B) Representative western blot of wild type and variant *SETD1B* protein in triplicates. (C) Western blot of GST-SET domain of *SETD1B* expressed in *E.coli* BL21, soluble fraction of protein extracts. (D) Representative melting curves of thermal shift analysis of GST-SET domain proteins. (F) Methylation variant pathogenicity scores for methylation profiles.

**Supplementary Figure S6:**
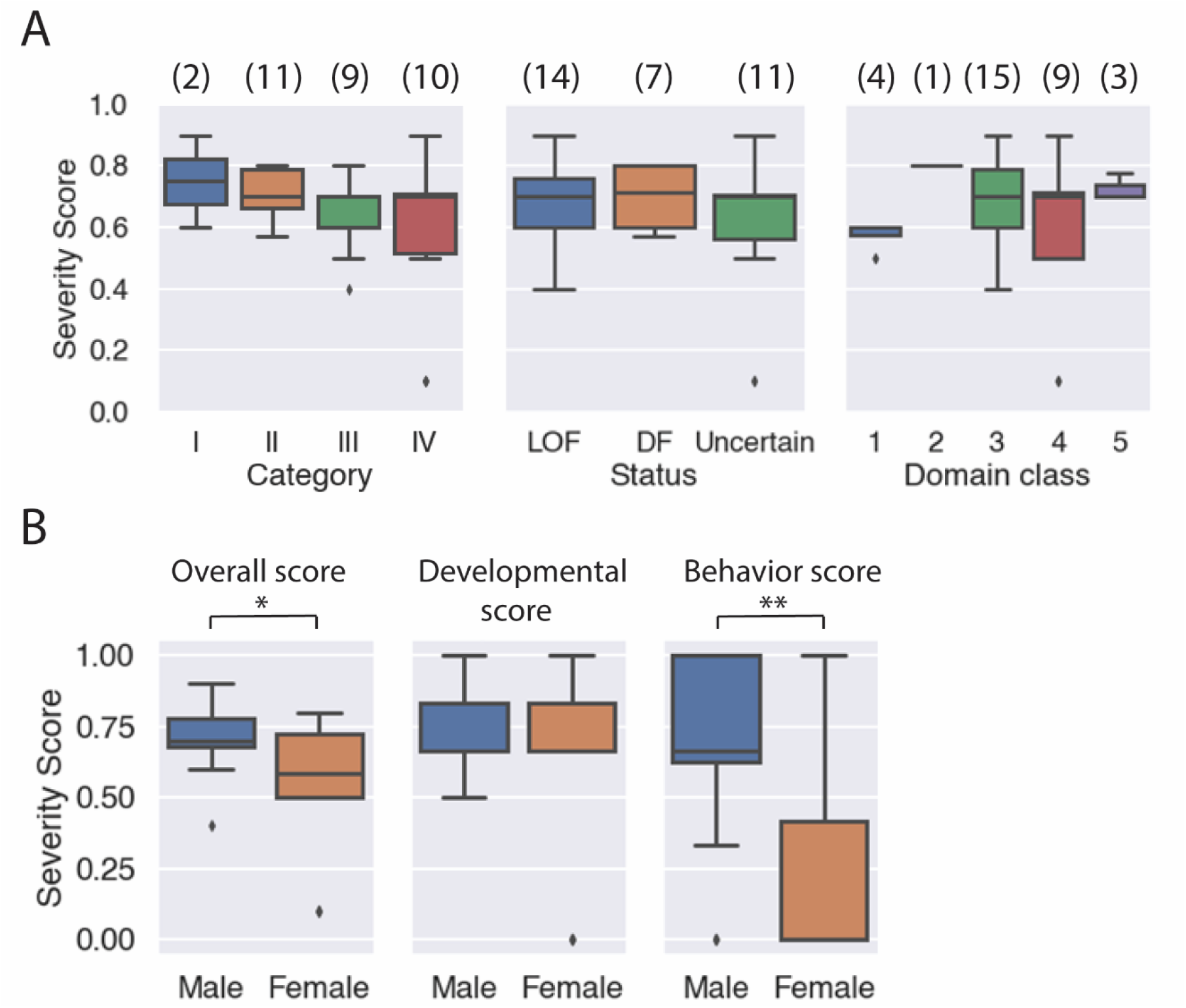
Average severity score in individuals with heterozygous variants. Severity scores were calculated for each individual, and the average for each category is shown only for heterozygous variants. A) severity score for categories for variants based on predicted effect on protein or stability (left plot, I=Catalytic site and/or substrate binding, II=Stability of *SETD1B* / complex formation / other protein-protein interactions, III=Truncation, IV=No apparent effect), functional status of the gene product (center plot, loss-of-function (LOF), diminished function (DF), or uncertain) or affected region (right plot, 1=All domains affected; 2=Middle, N-SET and SET domains affected; 3=N-SET and SET domains affected; 4=Middle region affected; 5=RRM domain affected). Numbers in parenthesis present the numbers of variants for each category. No statistically significant differences were observed (Welch Two Sample t-test p>0.05). B) heterozygous male (n=20) and heterozygous female (n=12) comparison for clinical features (left), only features related to development (center) or features related to behavior (right). The male and female groups are statistically different in the overall score (p=0.025) and in the behavior score (p=0.006) but not in the development score (p=0.5) (Welch Two Sample t-test).

## Supplementary Tables

**Supplementary Table S1:** Classification of all *SETD1B* variant (cohort and literature)

<<Supplementary Table S1.xlsx>>

**Supplementary Table S2:**
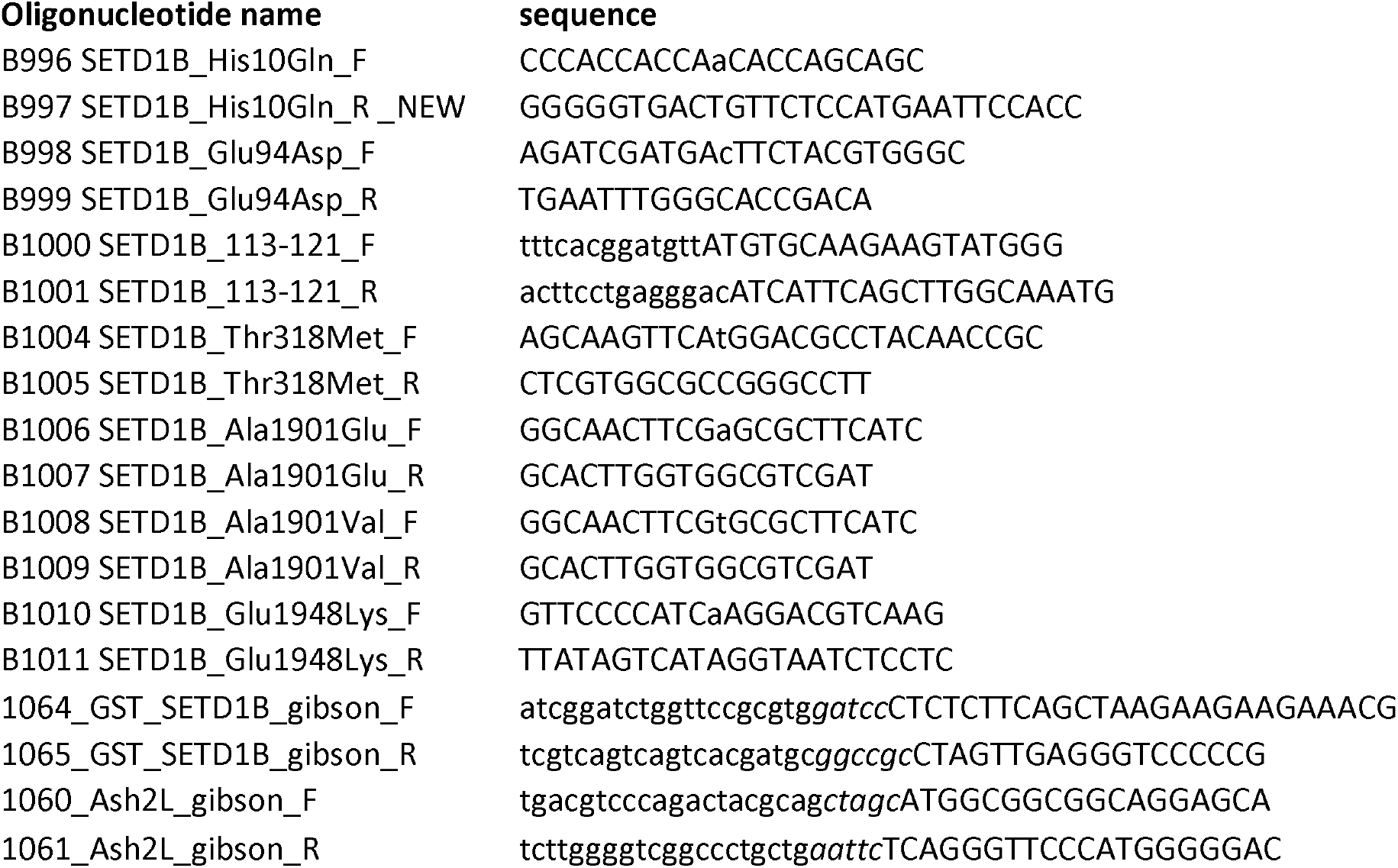
Oligonucleotides for site directed mutagenesis

**Supplementary Table S3:** Severity scores for all individuals in this study according to their clinical phenotype. Also annotated are functional effects, loss-of-function status, and the domains affected by each variant.

<<Supplementary Table S3.xlsx>>

